# Leveraging T-cell receptor – epitope recognition models to disentangle unique and cross-reactive T-cell response to SARS-CoV-2 during COVID-19 progression/resolution

**DOI:** 10.1101/2020.09.09.289355

**Authors:** Anna Postovskaya, Alexandra Vujkovic, Tessa de Block, Lida van Petersen, Maartje van Frankenhuijsen, Isabel Brosius, Emmanuel Bottieau, Christophe Van Dijck, Caroline Theunissen, Sabrina H. van Ierssel, Erika Vlieghe, Esther Bartholomeus, Wim Adriaensen, Guido Vanham, Benson Ogunjimi, Kris Laukens, Koen Vercauteren, Pieter Meysman

## Abstract

Despite the general agreement on the importance of T cells during SARS-CoV-2 infection, the clinical impact of specific and cross-reactive T-cell responses remains uncertain, while this knowledge may indicate how to adjust vaccines and maintain robust long-term protection against continuously emerging variants. To characterize CD8+ T-cell response to epitopes unique to SARS-CoV-2 (SC-unique) or shared with other coronaviruses (CoV-common), we trained a large number of TCR-epitope recognition models for MHC-I-presented SARS-CoV-2 epitopes from publicly available data. Applying those models to longitudinal COVID-19 TCR repertoires of critical and non-critical COVID-19 patients, we discovered that notwithstanding comparable CD8+ T-cell depletion and the sizes of putative CoV-common CD8+ TCR repertoires in all symptomatic patients at the initial stage of the disease, the temporal dynamics of putative SC2-unique TCRs differed depending on the disease severity. Only non-critical patients had developed large and diverse SC2-unique CD8+ T-cell response by the second week of the disease. Additionally, only this patient group demonstrated redundancy in CD8+ TCRs putatively recognizing unique and common SARS-CoV-2 epitopes. Our findings thus emphasize the role of the *de novo* CD8+ T-cell response and support the argument against the clinical benefit of pre-existing cross-reactive CD8+ T cells. Now, the analytical framework of this study can not only be employed to track specific and cross-reactive SARS-CoV-2 CD8+ T cells in any TCR repertoire but also be generalized to more epitopes and be employed for adaptive immune response assessment and monitoring to inform public health decisions.

## INTRODUCTION

The emergence of a severe acute respiratory syndrome coronavirus 2 (SARS-CoV-2) in 2019 has led to the most prominent pandemic in recent history. The SARS-CoV-2 infection manifests as coronavirus disease (COVID-19) with varying symptoms and severity and has caused substantial deaths all over the world.

The adaptive immune system is responsible for generating specific immunity against a viral infection. There is growing evidence that in case of SARS-CoV-2, T cells in particular might play a key role in infection control (Gangaev et al., 2021; Grifoni et al., 2020; Kared et al., 2021) even without seroconversion (Swadling et al., 2021; Wu et al., 2020), in moderation of COVID-19 severity (Chen and John Wherry, 2020; Tan et al., 2021), and in the durability of natural (Dan et al., 2021; Peng et al., 2020) and vaccination-induced (Afkhami et al., 2022; Tarke et al., 2022) immunity, including protection against viral variants. Previously, coronavirus-specific T-cell responses were already described as an important factor in the long-term immunity during SARS and MERS outbreaks (Liu et al., 2017; Oh et al., 2019; Yang et al., 2006), with some of the T cells exhibiting robust cross-reactivity against SARS-CoV-2 17 years later after the original infection (le Bert et al., 2020). Pre-infection presence of CD4+ (Braun et al., 2020; Mateus et al., 2020) and CD8+ (Lineburg et al., 2021; Nesterenko et al., 2021) T cells that broadly recognize epitopes of SARS-CoV-2 due to relatedness with previously encountered viruses have been widely reported. Furthermore, both cross-reactive CD4+ (Kundu et al., 2022; Loyal et al., 2021) and CD8+ (Mallajosyula et al., 2021; Schulien et al., 2021) T cells were suggested to have a protective effect in some individuals. Nevertheless, there are also conflicting findings questioning the clinical benefits of cross-reactive CD4+ (Bacher et al., 2020; Dykema et al., 2021; Saggau et al., 2022) and CD8+ (Ferretti et al., 2020) T cells for other patients. Consequently, it is important to investigate the clinical impact of pre-existing SARS-CoV-2-specific T cells and to be able to distinguish them from *de novo* responding T cells. In particular, understanding the contribution of CD8+ T-cell clones with a particular specificity could guide the design of new-generation vaccines or booster regimens to supplement weakening antibody-mediated neutralization (due to viral escape mutations) with an enhanced T-cell response and help stratify individuals into risk groups.

High-throughput T-cell receptor (TCR) sequencing paired with TCR-epitope mapping enables insights into an individual’s TCR repertoire composition. However, only a handful of TCR sequences, so-called “public” TCRs, are found across different individuals, while the majority of a TCR repertoire consists of “private”, unique to an individual, TCRs. Experimental assessment of epitope specificity of every single “private” TCR is not feasible due to the high inter- and intrapersonal diversity of TCRs. Accordingly, despite the ongoing efforts of sequencing studies to decipher “public” (Schultheiß et al., 2020; Snyder et al., 2020; Wu et al., 2022) and “private” (Minervina et al., 2021; Shomuradova et al., 2020; Snyder et al., 2020; Wu et al., 2022) SARS-CoV-2 TCR sequences, specificity of most disease-associated T cells is yet to be resolved.

Recently, computational recognition models have been developed to connect T cells with their target epitopes (Dash et al., 2017; de Neuter et al., 2017; Gielis et al., 2019; Glanville et al., 2017). These models are based on the concept that TCRs recognizing the same epitope tend to have similar amino acid sequences (Meysman et al., 2019). The advantage of the models is their ability to extract TCR-epitope interaction patterns from limited available experimentally validated data and to extrapolate them to previously unencountered TCRs. Thus, TCR repertoires can be easily screened to find potential epitope specificity of the unknown “private” and “public” TCRs. Here, we leverage such recognition models to track SARS-CoV-2 epitope-specificity in publicly available bulk TCR repertoires of COVID-19 patients (Schultheiß et al., 2020) and in sorted CD4+ and CD8+ TCR repertoires of newly recruited COVID-19 patients. We report on the differential evolution of CD8+ T-cell response to unique SARS-CoV-2 epitopes (SC2-unique) and SARS-CoV-2 epitopes that are shared with common cold coronaviruses (CoV-common) in patients with critical and non-critical COVID-19 presentation.

## MATERIALS AND METHODS

### SARS-CoV-2 epitope-TCR TCRex recognition models

#### • Collection of the public TCR data with known SARS-CoV-2 epitope-specificity

A collection of experimentally validated TCR-epitope pairs was established by combining two primary sources: (1) The VDJdb database which contained tetramer-derived data from Shomuradova et al (Shomuradova et al., 2020); access date: May 26th, 2020]; (2) The ImmuneCODE collection from Adaptive Technologies and Microsoft which contained pairs derived through MIRA assay (Snyder et al., 2020); access date: June 25th, 2020].

For all extracted pairs, several data curation steps were performed. All pairs matching more than one possible SARS-CoV-2 epitope were removed from the training data. Only valid TCR sequences that could be matched to standard IMGT were kept. To meet an internal TCRex limit, 5000 unique TCRs were selected randomly for epitope HTTDPSFLGRY.

#### • TCRex recognition model training and application

The paired TCR-epitope dataset was used to train a set of TCRex models using the standard procedures as described in (Gielis et al., 2019). In brief, for each epitope, TCRs experimentally validated to recognize the same epitope were used as a positive training data set to train a random forest model based on common physicochemical properties. Models were then evaluated using a 10-fold cross-validation, and any epitope-specific model with an AUC ROC higher than 0.7 and a PR higher than 0.35 was retained as suggested by the default TCRex settings. Models were constructed for all epitopes that had more than 30 distinct TCRs. All models used in this paper are available in the online TCRex tool (https://tcrex.biodatamining.be/).

TCRex models were then applied to the sequenced TCR repertoires of COVID-19 patients. Hits with a TCRex score greater than 0.9 and a baseline prediction rate (BPR) lower than 1e-4 were considered putative epitope-specific TCR sequences. Since the paired TCR-epitope dataset used to train TCRex models consists entirely of MHC class I restricted epitopes, valid predictions are only expected for CD8+ TCRs.

### Patient data

#### • “Split” dataset: patient cohort and separate TCR repertoire sequencing of presorted CD4+ and CD8+ T cells

The “split” dataset comprises separate CD4+ and CD8+ TCR repertoire sequences from patients with different COVID-19 severities collected during the initial stages of disease progression (the first 4 weeks after symptom onset). To this end, blood samples were collected from participants recruited in the IMSEQ study (NCT04368143), a prospective cohort study of COVID-19 patients admitted at the Antwerp University Hospital, Belgium (UZA) (the study was approved by the ITM IRB and UZA EC: number 20/12/135; ClinicalTrials.gov ID: NCT04368143). The inclusion criteria were: (1) having a laboratory-confirmed SARS-CoV-2 infection; (2) being older than 18; (3) providing written informed consent. COVID-19 severity was assigned based on the worst symptoms observed during the entire course of the disease.

For TCR sequencing of the COVID-19 patients, individuals exceeding the age of 65 and individuals diagnosed with or treated for oncologic conditions were excluded. We further selected patients that had donated at least two consecutive blood samples taken at least two days apart, the first of which within 16 days of symptom onset. As a result, 11 individuals with confirmed SARS-CoV-2 infections were retained including 7 patients classified as moderate, 1 as severe, and 3 as critical according to WHO grading (World Health Organization, 2021) summarized in Table S1. Characteristics and sampling time points of retained study volunteers are summarized in Table S2.

At each time point, whole blood samples were obtained using three 9 mL S-Monovette® lithium heparin tubes (Sarstedt). The PBMC fraction was isolated using LymphoprepTM (StemCell technologies), before cryopreserving aliquots in liquid nitrogen until further use.

After thawing, CD4+ and CD8+ T cells were positively selected using magnetic MicroBeads (Miltenyi Biotec), as described by the manufacturer. Counting was done manually on Trypan blue-stained cells using C-Chip counting chambers (NanoEnTek). All samples contained at least 200.000 viable cells and were stored in DNA/RNA shield (Zymo) at −80°C. Total RNA was extracted using Quick RNA microprep kit (Zymo) following the manufacturer’s protocol, eluted in 18μl DNAse/RNAse free H2O. The RNA concentrations were determined with Qubit RNA HS assay kit (Thermo Fisher Scientific). Each sample was split into triplicates (i.e., 3 vials of 5μl) that were used as library prep input. TCR library prep was done with QIAseq® Immune Repertoire RNA Library and QIAseq®index kit (Qiagen, Venlo, Netherlands) that amplifies TCR alpha, beta, gamma, and delta chains. After quality control using TapeStation (Agilent, Santa Clara, CA, USA), concentration was measured with the Qubit dsDNA HS Assay kit (Thermo Fisher Scientific). For sequencing, each library was equivolume pooled. The pool was diluted to 4 nM and denatured. 1.1 pM of denatured library pool was run on the NextSeq 500 (NextSeq 500/550 Mid Output Kit v2.5, Illumina Netherlands) using 300 cycles with a pair-end 261-8-8-41 base read.

#### • “Mixed” dataset: collection of the publicly available bulk TCR repertoire sequences (CD4+ and CD8+ together)

A complementary “mixed” dataset (Schultheiß et al., 2020), which contains TCR repertoires sequenced in bulk, without prior T-cell sorting into CD4+/CD8+, was downloaded from the iReceptor gateway (Corrie et al., 2018) [access date: July 13th, 2020]. It features longitudinal (from week 1 to 8 after symptom onset) samples taken from patients with active disease and single time point samples from those that have recovered (week 4+ after symptom onset), summarized in Table S2. Patients classified as “severe” in the original study were reclassified as “critical” to match WHO grading (World Health Organization 2021) (Table S1). The data of asymptomatic individuals were excluded as their TCR response lies outside of the scope of our study. In total, TCR repertoires of 36 individuals were included (Table S2): 24 non-critical patients (16 recovered individuals who had mild COVID-19, 7 active patients with moderate COVID-19 and 1 patient with mild COVID-19 who had data points from both active and recovery phase of the disease) and 11 active patients with critical COVID-19 presentation. Noteworthy, critical disease resulted in the death of 5 out of 11 critical patients.

#### • Merged dataset: CD8+ and CD4+ TCR repertoires of the “split” dataset together with CD8+ TCR repertoires of the “mixed” dataset

Patient data analyzed in this study were assembled by combining TCR repertoires from internally produced “split” and publicly available “mixed” datasets into one merged dataset of 46 TCR repertoires of symptomatic COVID-19 patients. To make the number of data points per week more comparable between disease severity groups (Fig.S1-a, b), all individuals were divided into two groups (Table 1): critically ill (14 patients with critical COVID-19 severity) and non-critically ill (32 patients with mild, moderate, or severe COVID-19 severity). All available data points from the “split” dataset (11 symptomatic patients) were also used separately to verify whether the predictions of constructed recognition models are specific for the CD8+ T-cell subset.

**Table 1.**
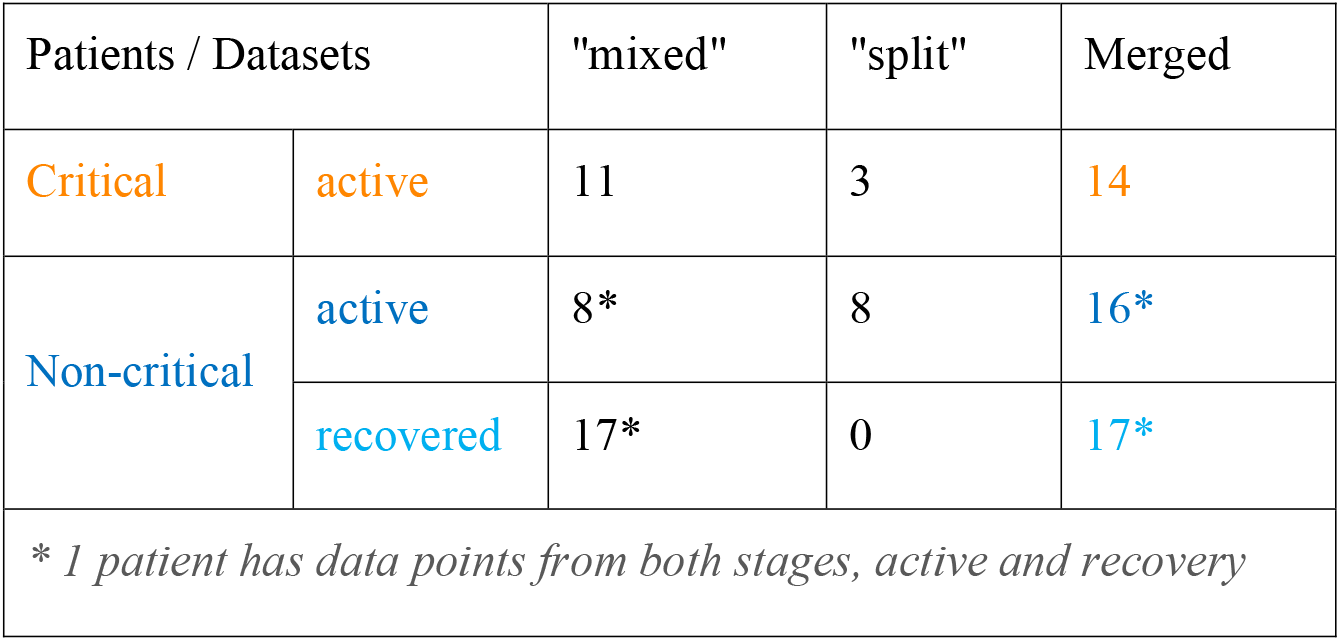
Summary of the datasets used for the analysis in this study

| Patients / Datasets |  | "mixed" | "split" | Merged |
| --- | --- | --- | --- | --- |
| Critical | active | 11 | 3 | 14 |
| Non-critical | active | 8* | 8 | 16* |
|  | recovered | 17* | 0 | 17* |
| * 1 patient has data points from both stages, active and recovery |  |  |  |  |

To compensate for shifts in the response onset due to diverse times of admission and differences in the number of available data points for each patient, in further analysis of TCR repertoire metrics, only the maximum values of each week were considered when multiple time points were available for the same person during every time interval, (i.e., maximum TCR fraction between days 1-7 for week 1, maximum number of recognized epitopes between days 8-14 for week 2, etc.). Consequently, out of 32 individuals in the non-critical patient group, 15 patients had data points from the active stage of COVID-19, 16 – from the recovery stage, and 1 – from both stages (Table 1). All 14 critically ill patients remained sick throughout the entire duration of the study (up to 8 weeks) (Table 1).

### TCR repertoire data processing and analysis

Demultiplexing of the sequencing data, UMI correction and generation of the UMI consensus for the “split” dataset were performed using MiNNN v.10.1 (https://minnn.readthedocs.io/). As three technical replicate experiments were conducted for each sample, only those TCR sequences that occurred in at least two out of three replicates were kept. Out of the selected replicates, the one with the highest total TCR count was retained for the downstream analysis.

Further steps were identical for both “split” and “mixed” datasets. TRB (T cell receptor beta gene) clonotype annotation was performed using MiXCR v. 3.0.13 with the default input parameters (Bolotin et al., 2015). Only those TCRs that occurred at a frequency of at least 1 in 100 000 were retained, to compensate for the different sequencing depths between studies. Metadata was made uniform so that the time points are annotated by weeks after the onset of symptoms. All the data processing, comparisons and statistical analysis were performed using standard python3 libraries. Code necessary to enable the reproduction of the processing and analysis steps can be found in GitHub repository [https://github.com/apostovskaya/CovidTCRs/src].

### SARS-CoV-2 epitopes

To establish the “uniqueness” of each epitope in our database, we compared the presence of every SARS-CoV-2 epitope against a list of protein sequence data for 119 Nidovirales species, including SARS-CoV and human common-cold coronaviruses. These data were retrieved from the Corona OMA Orthology Database (Altenhoff et al., 2018), where the used protein amino acid sequences for SARS-CoV-2 correspond to Genbank accession GCA_009858895.3, and the protein amino acid sequences for SARS-CoV to GCA_000864885.1. SARS-CoV-2 epitopes were matched to all proteins of all 119 species with an exact match, as the degree of variation allowed in the epitope space while retaining TCR recognition is still an unsolved question. Sequence identity between proteins was established using a pairwise protein BLAST. Matches across all species for each epitope were tallied, and the annotation for SARS-CoV-2 was retained: SARS-CoV-2 epitopes that occur only in 1 species (SARS-CoV-2) were labeled as “SARS-CoV-2-unique” (SC2-unique) and all others as “common for coronaviruses” (CoV-common).

### T-cell receptor metrics

Different approaches can be used to analyze TCR repertoires. In our case, we were specifically interested in CD8+ T cells recognizing SARS-CoV-2 epitopes. Thus, four parameters were selected as the most informative: CD8+ TCR repertoire depth, CD8+ TCR repertoire breadth, CD8+ T-cell response diversity and CD8+ T-cell response redundancy. Repertoire depth was described as the relative frequency with which TCR sequences with a certain predicted specificity occur in the entire TCR repertoire. Repertoire breadth was calculated as the number of unique TCR sequences with a certain predicted specificity divided by the size of the unique TCR repertoire. Response diversity was represented as the number of putatively recognized SARS-CoV-2 epitopes. Average response redundancy was estimated as TCR/Epitope ratio: the number of unique SARS-CoV-2-specific TCRs to the number of recognized SARS-CoV-2 epitopes. Response metrics were calculated separately for SC2-unique and CoV-common epitopes. Repertoire metrics were rescaled so that proportions are consistent across data sets. Additionally, log2 fold change of a repertoire depth was monitored to evaluate the magnitude of temporal intrapersonal changes. Accordingly, for every patient for whom longitudinal data was available, TCR repertoire frequencies were shifted by 1 to enable subsequent calculation of fold changes between consecutive weeks and log2-transformation.

## RESULTS

### SARS-CoV-2 epitope-TCR recognition models are robust and performant

To construct SARS-CoV-2 epitope-TCR recognition models, a collection of experimentally validated TCR-epitope pairs was established. To this end, data derived from T cells, the specificity of which was identified with peptide-MHC tetramers (Shomuradova et al., 2020) was combined with ImmuneCODE TCR-epitope pairs derived from sorting of antigen-stimulated and activated CD8+ T cells using MIRA (multiplex identification of TCR antigen specificity) assay (Snyder et al., 2020). After curation of the data and quality filtering of the models, 47 distinct epitope TCRex models were retained for SARS-CoV-2. An overview of all models and their performance can be found in Table S3.

The number of newly constructed TCRex models for SARS-CoV-2 epitopes almost equals 49 previously available TCRex models for all non-SARS-CoV-2 epitopes combined (Gielis et al., 2019), indicating the vast amount of data that has been generated since the start of the pandemic compared to what has been collected for all prior pathogens and diseases. Twenty-four of these 47 epitopes match the SARS-CoV-2 replicase protein coded by ORF1ab, 16 match the SARS-CoV-2 spike protein encoded by ORF2 and the final 7 are distributed across the remaining proteins (Fig.1). In addition, 19 of the 47 epitopes are 100% unique to SARS-CoV-2 in our dataset of 119 Nidovirales species. As can be seen in figure 1, the unique SARS-CoV-2 epitopes are not evenly distributed across the proteins. Whereas only 6 out of 24 are unique for the ORF1ab replicase protein, 9 out of the 16 epitopes derived from the spike protein are unique to SARS-CoV-2. Although mutations are 5 times more frequent in the spike protein compared to the genomic average (Amicone et al., 2022) and thus might weaken B-cell response, T-cell recognition appears to be retained against different variants including Omicron (Moss, 2022; Redd et al., 2022; Tarke et al., 2022). Additionally, as previously evaluated for 3 SARS-CoV-2 variants, only 1 mutation overlapped with 52 MHC-I epitopes recognized by convalescent individuals (Redd et al., 2021). Together, this bodes well for the constructed TCRex models to stay relevant for the continuously emerging variants.

**Figure 1.**
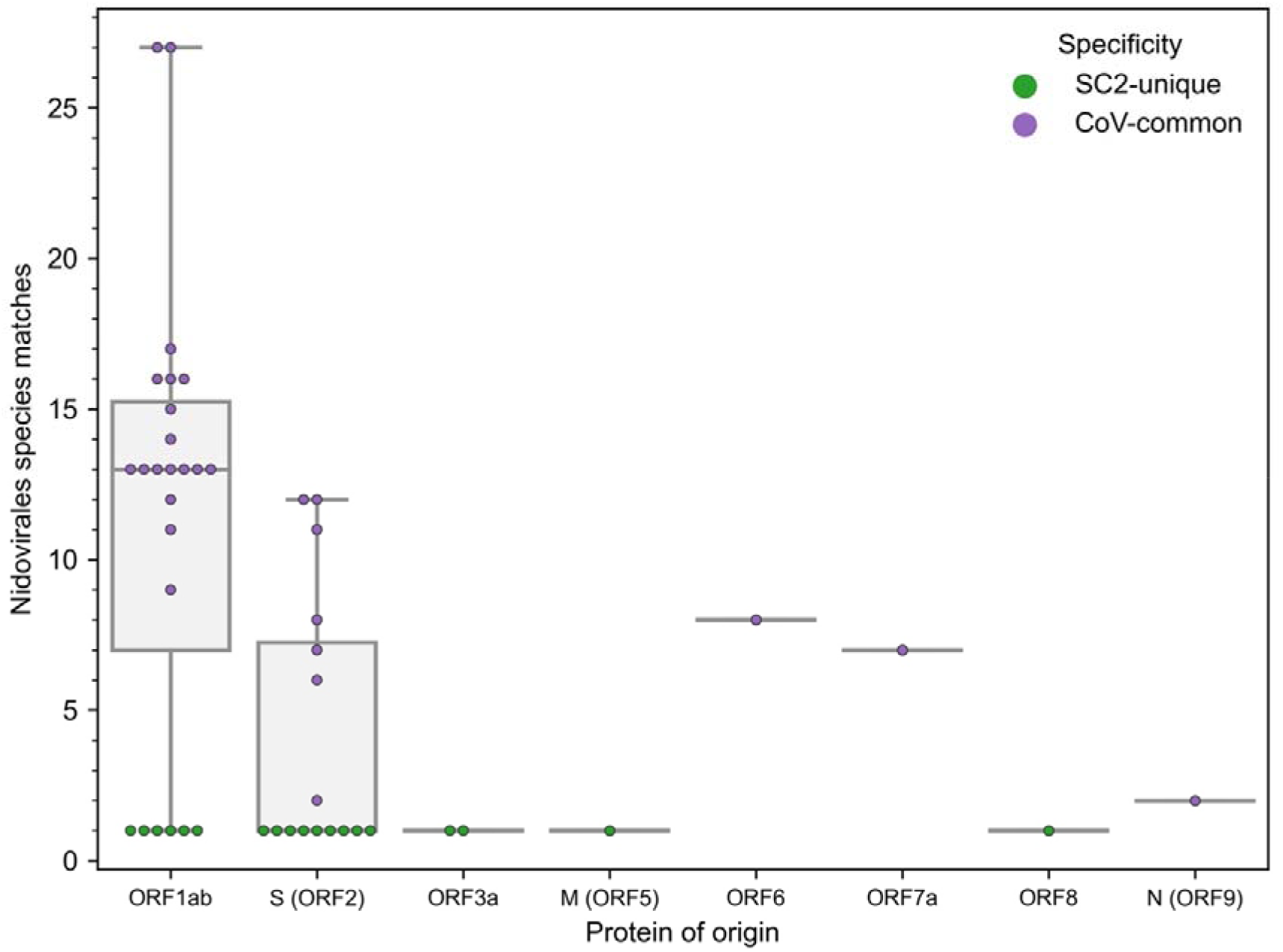
Distribution of the 47 epitopes for which TCR recognition models could be created across SARS-CoV-2 proteins (x-axis) and amount of exact amino acid epitope matches within the 119 species of the Nidovirales order (y-axis). TCR models for epitopes unique to SARS-CoV-2 (n=19) correspond to y=1.

As we were integrating models from different resources and diverse experimental methods, we wished to confirm if these data were comparable. Interestingly, one epitope had both tetramer (315 TCRs) and MIRA data (366 TCRs), namely YLQPRTFLL (YLQ). However, in the case of the MIRA data, the TCRs were not uniquely assigned to this epitope, but all were assigned to the trio YLQPRTFL, YLQPRTFLL, and YYVGYLQPRTF. These 366 TCRs were thus excluded from the training data. Therefore, MIRA YLQ data can serve as an independent dataset for performance evaluation of the YLQ model based on tetramer YLQ data. In this manner, TCRex predicted 81 out of 366 TCRs in the YLQ MIRA dataset to be putatively YLQ-reactive. Notably, only 35 out of 81 TCRs had an exact sequence match between the two datasets based on CDR3 sequence (not accounting for V/J genes); hence, 46 additional TCRs were identified, indicating TCRex can indeed extrapolate from found TCR patterns. This number of predictions was assigned an enrichment p-value of 6.44e-246 based on the built-in TCRex enrichment test. As a negative control, no other epitopes present in TCRex (including both the 46 non-YLQ SARS-CoV-2 models and the 49 non-SARS-CoV-2 models) were predicted to have a single TCR target within the YLQ MIRA dataset. Thus, the data is comparable, and the models can be used irrespective of origin.

### Recognition models can be used to track epitope-specificity in CD8+ TCR repertoires

As the constructed models predict specificity to epitopes presented on MHC-I molecules, we expected that TCRs predicted to recognize those epitopes will be enriched in CD8+ T cells. To validate this assumption, we generated CD4+ and CD8+ TCR repertoires for 11 COVID-19 patients (“split” dataset) at multiple time points and applied our 47 SARS-CoV-2 TCRex models to this data. As can be seen in Figure 2, the number of predicted SARS-CoV-2 reactive TCRs was indeed significantly higher in the CD8+ compared to the CD4+ T-cell population (MU, p=0.001). There is, however, a small number of hits within the CD4+ population which is not unforeseen given some inefficiency inherent to magnetic cell sorting and common CDR3 sequences occurring in both the CD4+ and CD8+ populations (Carter et al., 2019; Meysman et al., 2019). The predominant signal in the CD8+ T cells confirms that the models are specific towards this subpopulation and thus suitable to track particularly the CD8+ T-cell response in any individual TCR repertoire, sequenced in bulk or after prior sorting.

**Figure 2.**
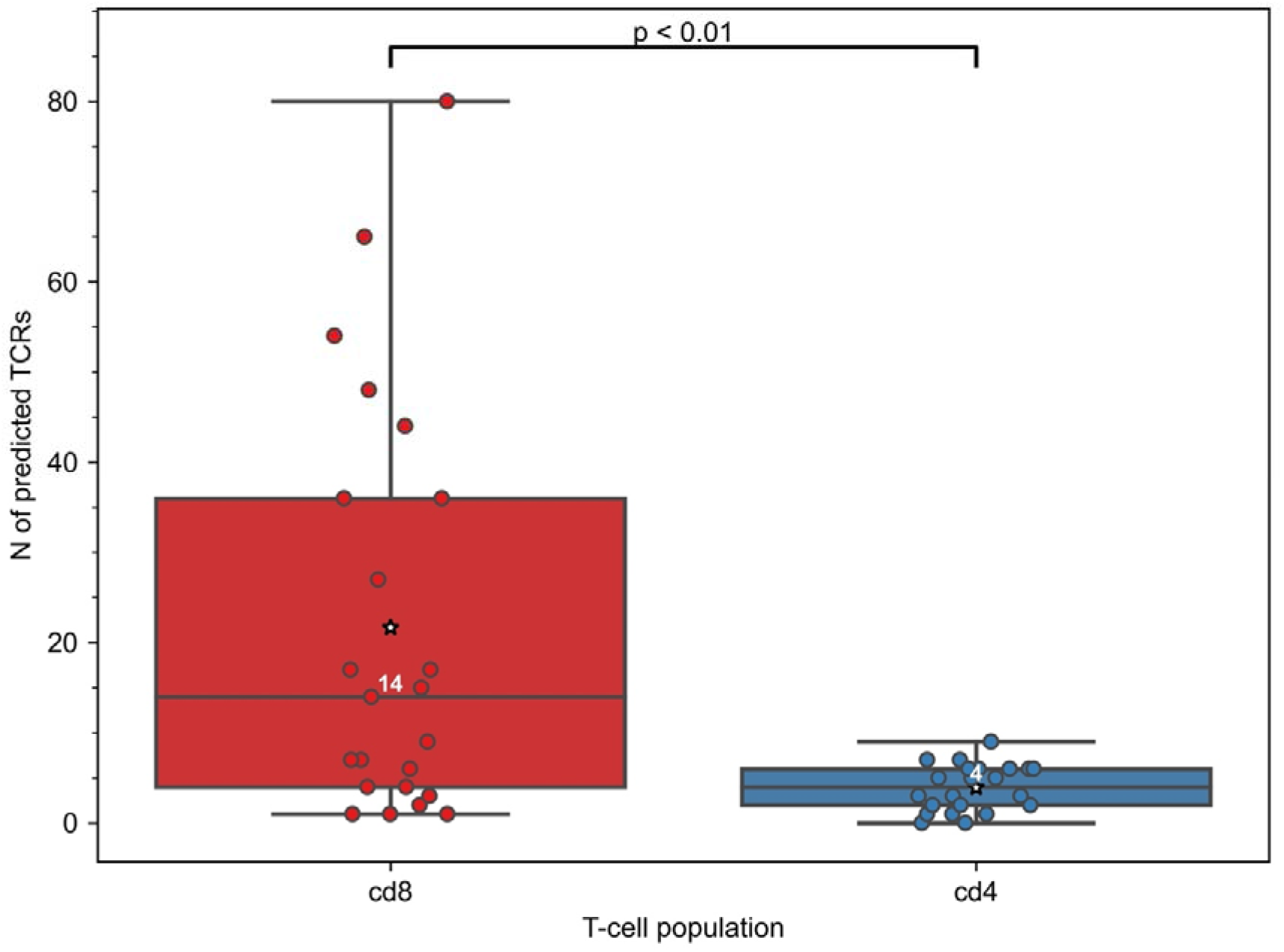
Constructed TCRex recognition models are suitable for the prediction of CD8+ T-cell specificity. As the models were built for epitopes presented in MHC-I, the number of TCRs predicted to recognize SARS-CoV-2 epitopes was significantly higher in CD8+ than in CD4+ T-cell repertoires when SARS-CoV-2 TCRex models were applied to an in-house COVID-19 patient “split” TCR dataset (MU, p=0.001, nCD8=23, nCD4=22). White numbers specify the median number of the TCRs in a repertoire that were predicted by TCRex to be specific to SARS-CoV-2 epitopes; mean values are represented by a star.

### Initial CD8+ T-cell response is similar in all patients, regardless of COVID-19 severity

In this study, we employed TCRex models to analyze COVID-19 TCR repertoires of 14 critically and 32 non-critically ill symptomatic patients. Prediction of putative SARS-CoV-2-specific CD8+ T cells identified 755 TCRs in the dataset cohort. Of these, 149 and 606 TCRs were found in samples from patients with critical and non-critical COVID-19 presentation, respectively.

Since the level of pre-existing T cells cross-reactive to SARS-CoV-2 and the swift mounting of T-cell responses had been postulated to influence COVID-19 progression (Sette and Crotty, 2021; Tan et al., 2021), we first assessed the initial size of SARS-CoV-2 specific CD8+ TCR repertoires. Therefore, the differences in the abundance of T-cell clones putatively recognizing MHC-I presented SARS-CoV-2 epitopes that are either unique to the virus (SC2-unique) or also occur in other *Nidovirales* species (CoV-common) were compared in patients with critical and non-critical symptomatic COVID-19. The prevalence of CD8+ TCRs with a certain specificity was described as relative frequencies with which those TCRs occur in the collected TCR repertoires, i.e., the depth of the responding TCR repertoire.

All active patients, regardless of the disease severity, had more TCRs specific to CoV-common than SC2-unique epitopes only during week 1 after the symptom onset (Fig.3) and not at any other subsequent week of the disease (Table S4). This disparity was more pronounced in critical patients for whom the difference was statistically significant (Bonferroni corrected Mann–Whitney U test p=0.048, AUC=1, Fig.3B). In the non-critical group, the frequency of putative SC2-unique TCRs was already higher than in the critical group, although not significantly (Table S5). In addition, no significant difference in the frequency of CoV-common TCRs, the total number of TCRs and the percent of unique TCRs was detected between critical and non-critical groups at this time (Table S5). These findings suggest that prior to the encounter with SARS-CoV-2, many individuals already have a substantial TCR repertoire dedicated to CoV-common epitopes.

**Figure 3.**
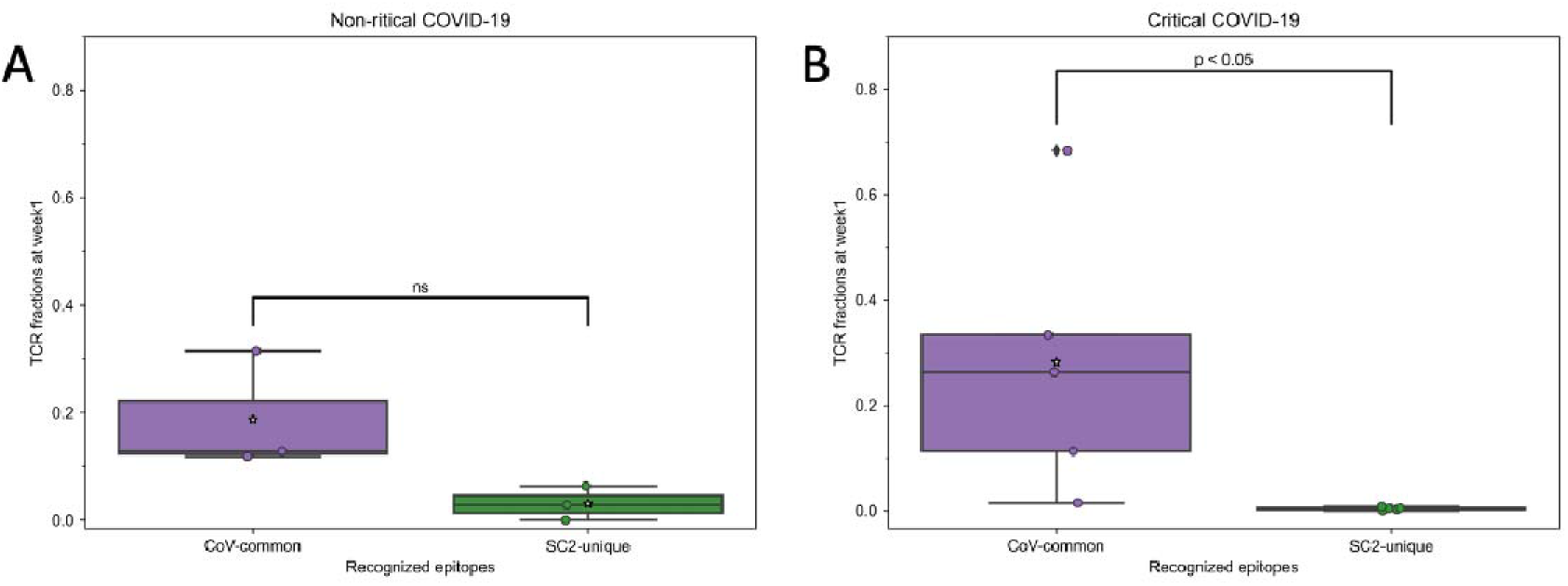
During the first week of COVID-19, relative frequencies of TCRs (depth of the repertoire) predicted to recognize CoV-common epitopes were higher than of TCRs putatively specific to SC2-unique epitopes in all patients with (**A**) non-critical (Bonferroni corrected Mann–Whitney U test p=0.16, n=3) and (B) critical (Bonferroni corrected Mann–Whitney U test p=0.048, n=5) COVID-19. Mean values are represented by a white star.

### Putative SARS-CoV-2 specific CD8+ T cells are mounted within the first two weeks of COVID-19 only in non-critical patients

To investigate whether the differences between frequencies of SC2-unique and CoV-common TCRs ceased after week 1 due to the increase of SC2-unique or the decrease of CoV-common TCR repertoire, we studied changes in the depth (frequency of specific TCRs) and breadth (percent of specific TCRs out of unique TCRs) of respective TCR repertoires within each patient group. We observed that critical and non-critical patients had the opposite dynamics during the first two weeks (summarized in Table 2): the median breadth and depth of SC2-unique TCR repertoires increased from week 1 to week 2 only in non-critical patients (Fig.S2-a, c), and the median breadth and depth of the CoV-common TCR repertoire decreased from week 1 to week 2 only in critical patients (Fig.S2-b, d). Furthermore, we compared intragroup changes in the diversity of the response – the number of different SC2-unique and CoV-common epitopes being recognized by an individual TCR repertoire. We reasoned that an increase in those parameters could be an indirect indication of the *de novo* activation of T cells as opposed to the expansion of already activated T-cell clones. We discovered that non-critical patients, unlike critical ones, generally recognize 2 times more CoV-common and 3 times more SC2-unique epitopes at week 2 compared to week 1 (Table 2, Fig.S2-e, f). Additionally, only the redundancy of SC2-unique response increased 10 times between weeks 1 and 2 in non-critical patients alone (Table 2, Fig.S2-g, h).

**Table 2.** Changes in the SARS-CoV-2 TCR repertoire between week 1 and week 2.

| Specificity | Parameter |  | Patients |  |
| --- | --- | --- | --- | --- |
|  |  |  | Critical | Non-critical |
| CoV-common | TCR repertoire | depth (TCR frequency) | ↓ | = |
|  |  | breadth (% of specific TCRs) | ↓ | = |
|  | Response | diversity (No. recognized epitopes) | ↓ | ↑ |
|  |  | redundancy (No. unique TCRs per 1 epitope) | = | = |
| SC2-unique | TCR repertoire | depth (TCR frequency) | = (0) | ↑ |
|  |  | breadth (% of specific TCRs) | ↓ | ↑ |
|  | Response | diversity (No. recognized epitopes) | ↓ | ↑ |
|  |  | redundancy<br>(No. unique TCRs<br>per 1 epitope) | = | ↑ |

All these results consistently point out that even though the CD8+ T-cell response in all symptomatic patients, regardless of the disease severity, starts with the mounting of (pre-existing) T cells specific to CoV-common epitopes during the first week after the symptom onset, only individuals with the non-critical disease appear to be effectively activating and expanding T cells recognizing SC2-unique epitopes during the first two weeks of the disease. Intriguingly, we have observed a lack of increase in CoV-common TCR repertoire depth and breadth and CoV-common response diversity in non-critical patients, despite the growth in the number of recognized epitopes. This could be attributed to the two opposing processes balancing each other out: continuous depletion and *de novo* activation of T cells recognizing CoV-common epitopes.

### COVID-19 severity is moderated by SC2-unique TCR repertoire depth, redundancy of SC2-unique and diversity of CoV-common TCR responses

Since the development of CD8+ T-cell response to SARS-CoV-2 differed during the first two weeks depending on the disease severity, we further compared the response levels between 7 critically and 7 non-critically ill patients at week 2, when TCRs to both previously seen and newly encountered epitopes are expected to have been activated and expanded. Our results revealed that in 81.6% of pairwise comparisons between individuals from different disease severity groups, non-critical patients had significantly higher frequencies of TCRs specific to SC2-unique (Bonferroni corrected Mann–Whitney U test p=0.039, Fig.4A) but not CoV-common (Bonferroni corrected Mann–Whitney U test p=0.456, Fig.4B) epitopes than critically ill patients during week 2 of COVID-19. Moreover, we observed that although the total number of TCRs differed significantly between the groups based on patient disease severity at week 2 (Bonferroni corrected Mann–Whitney U test p=0.033, AUC=0.878, Fig.S3-a) and not at week 1 (Table S5), the proportion of unique TCRs in repertoires did not (Fig.S3-b).

**Figure 4.**
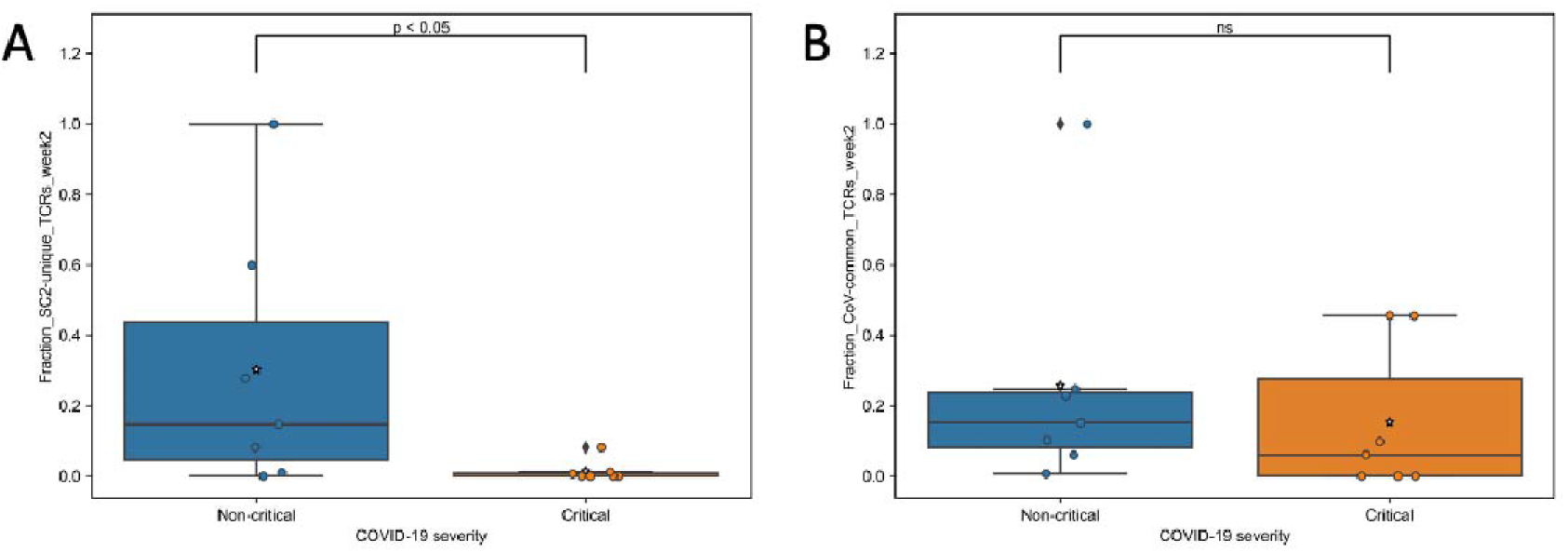
During the second week of COVID-19, relative frequencies of TCRs (depth of the repertoire) predicted to recognize (**A**) SC2-unique (Bonferroni corrected Mann–Whitney U test p=0.039, n=14 (7 critical and 7 non-critical)) but not (**B**) CoV-common (Bonferroni corrected Mann–Whitney U test p=0.456, n=14 (7 critical and 7 non-critical)) epitopes were significantly higher in symptomatic non-critical compared to critically ill patients. Mean values are represented by a white star.

To gather more insights into whether the increase in SARS-CoV-2 specific TCRs is caused by expanding the amount of T cells recognizing a limited number of epitopes or by increasing the number of epitopes being recognized by the repertoire, we compared the overall diversity of the response in our symptomatic patient groups. The number of recognized CoV-common but not SC2-unique epitopes was significantly higher at week 2 in the non-critical patient group (Bonferroni corrected Mann-Whitney U test p=0.026, AUC=0.796, Table S5). When this parameter was normalized for the repertoire size, there was no difference (Table S5) suggesting that the CD8+ T-cell response of critical patients was limited by the low number of available unique TCRs (diminished TCR diversity). Furthermore, non-critical patients, unlike critical patients (Fig.5A,B, median=1 for both epitope groups), exhibited redundancy in the T-cell response to both SC2-unique (Fig.5A, median=10 TCRs “per average epitope”, range=[1-26]) and, although much less pronounced, CoV-common (Fig.5B, median=2 TCRs “per average epitope”, range=[1-3]) epitopes. Later (week 3+), the redundancy of the response to both groups of epitopes was decreasing in active non-critical patients (Fig.S4) and disappeared after patients had recovered (Fig.5A,B). During these later weeks, active critical patients also started to develop redundancy but only to SC2-unique epitopes (Fig.S4).

**Figure 5.**
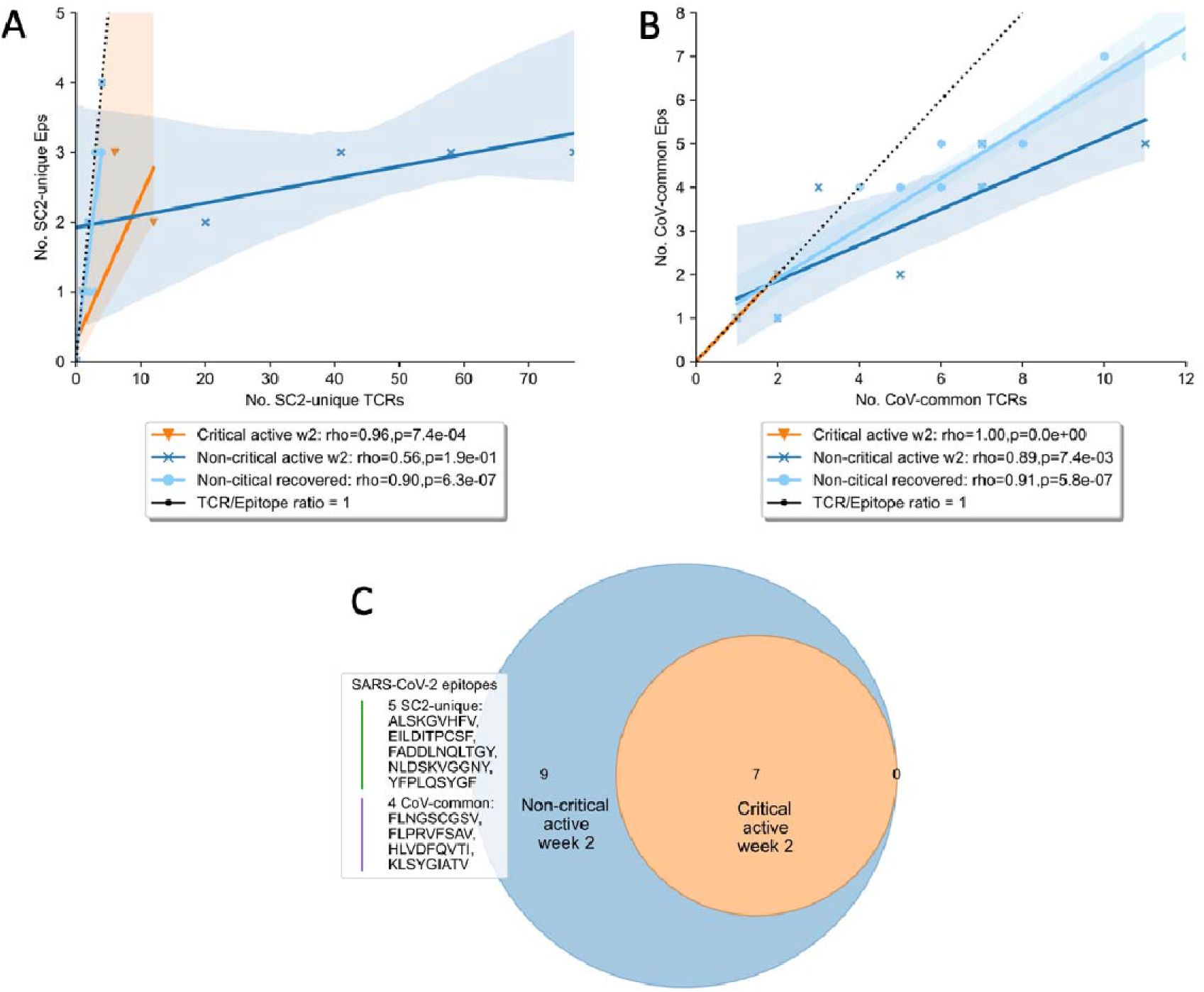
Diversity and redundancy of the response differed between patients with critical and non-critical symptomatic COVID-19 during the second week of the disease. (**A, B**) Redundancy (the number of unique TCRs recognizing the same epitopes) was significantly higher in non-critical (dark blue) compared to critical (orange) patients and disappeared once patients recovered (light blue). This was true for both (**A**) SC2-unique and (B) CoV-common epitopes. (**C**) 9 epitopes, out of which 5 are unique to SARS-CoV-2 (SC2-unique) and 4 are shared with other species of the Nidovirales order (CoV-common), were recognized exclusively by non-critical patients during the second week of COVID-19. No epitopes were recognized only by critical patients.

Since at this time all patient groups had TCRs putatively targeting both SC2-unique and CoV-common epitopes, we also set out to investigate whether there was a specific set of epitopes recognition of which might lead to a milder case of COVID-19. Unsurprisingly, no epitopes were recognized by all the patients within any of the two disease severity groups. As epitope recognition depends not only on the TCR sequences but also on what epitopes can be presented given the HLA type that a person has, it is most likely that sets of recognized epitopes are unique for each patient. However, there were 9 epitopes, including 5 SC2-unique ones, that were recognized exclusively by non-critical patients, albeit by only 1 or 2 individuals for each epitope (Fig.5C). Those epitopes were recognized throughout the entire duration of the disease, by active or recovered non-critical patients.

Together, our results indicate that pre-existing SARS-CoV-2 cross-reactive CD8+ T cells alone could not be specifically related to symptomatic non-critical COVID-19. Critically ill patients struggle to mount SC2-unique CD8+ TCRs and sustain CoV-common CD8+ TCRs while in non-critical patients, CD8+ T-cells putatively recognizing SARS-CoV-2, and especially those targeting SC2-unique epitopes, are activated and expanded by the second week of COVID-19. Less severe COVID-19 also seems to be associated with a diverse and redundant CD8+ T-cell response rather than recognition of specific epitopes.

### TCR diversity potential is reduced during COVID-19

To gain a better understanding of the SARS-CoV-2 associated changes in the overall TCR repertoire, we next analyzed longitudinal data spanning 8 weeks. As reported above, critical and non-critical symptomatic patients had comparable total numbers of sampled TCRs and proportions of unique TCRs at the beginning of their COVID-19. However, examination of all available data revealed that the proportion of all unique TCRs (irrespective of their epitope specificity) significantly increased in individuals with the non-critical disease over the entire period of the study (Fig.6, Spearman rho=0.62, p=3e-05) and became significantly higher after week 4, once patients entered recovery stage (Mann–Whitney U test p=2e-04, AUC=0.83). In contrast, TCR repertoires of critical patients, despite some trend for increase between weeks 3 and 6, on average remained less diverse even at week 8 of the disease (Fig.6, rho=0.08, p=0.69) indirectly supporting a previously reported correlation between disease severity and lymphopenia derived from a different type of analysis (Tan et al., 2020). Those findings suggest that both patient groups experience similar depletion of CD8+ T cells during the early phase of the SARS-CoV-2 infection, and this depletion is preserved in active critical patients but restored in recovered non-critical patients.

**Figure 6.**
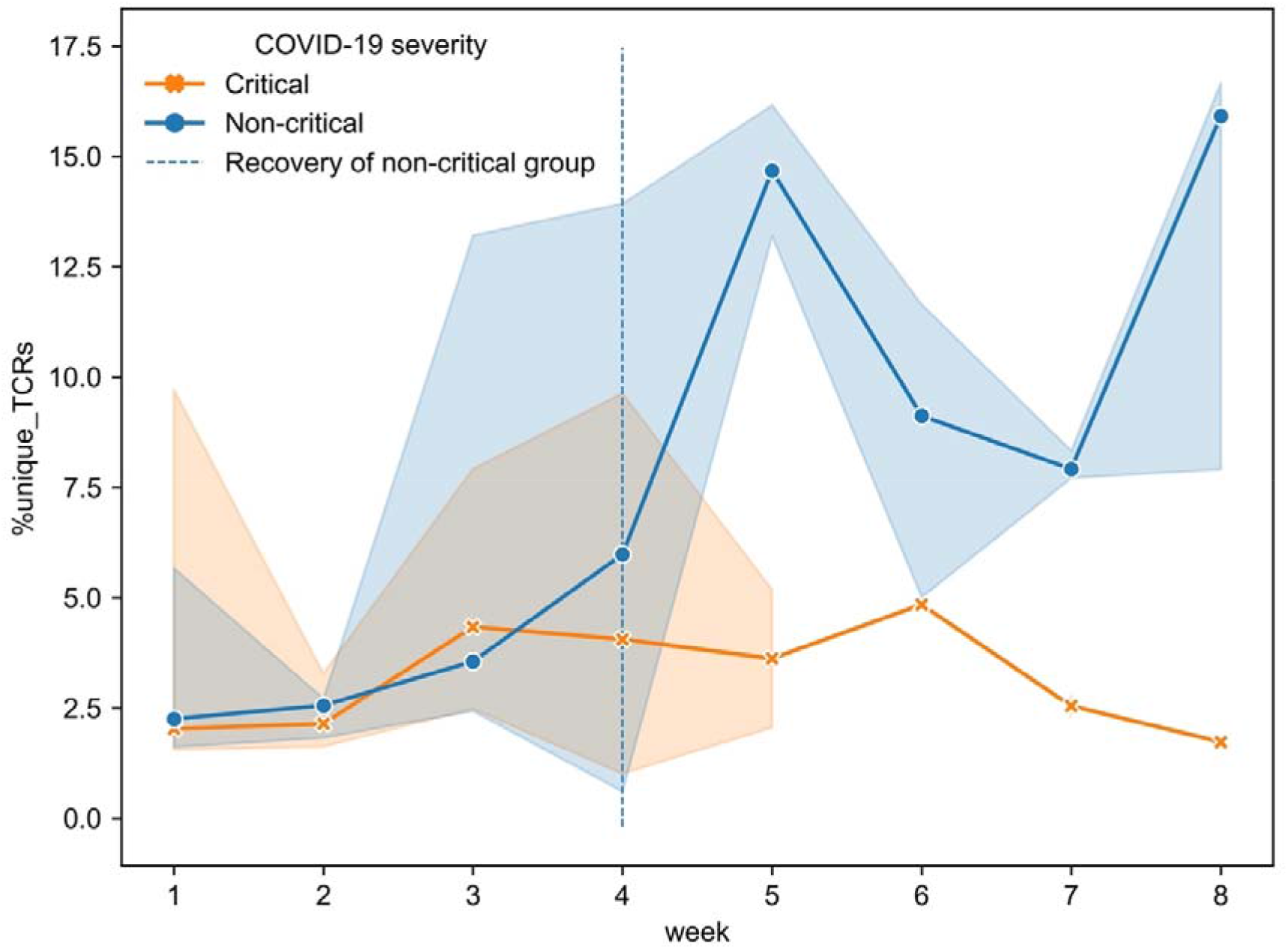
Proportion of unique TCRs was increasing significantly only in symptomatic non-critically ill patients (dark blue, Spearman rho=0.62, p=3e-05) and became significantly higher once patients started recovering (Mann–Whitney U test p=2e-04). Multiple inter- and intra-individual values combined within each disease severity group (n = 14 critical, 32 non-critical) are represented as tendency lines with a 95% confidence interval when multiple data points were available at overlapping time points, shown as shadow areas. The vertical dashed line (black) separates the active and recovery stages of the disease in non-critical patients.

### Development of SARS-CoV-2 reactive CD8+ T-cell immunity in critical patients is dominated by TCRs predicted to target epitopes unique to SARS-CoV-2

To further investigate the most prominent dynamics of SARS-CoV-2 TCR repertoires, we disentangled longitudinal changes in SC2-unique and CoV-common TCRs in our patient cohorts with different COVID-19 severity. First, we considered only patients of whom samples were available from at least two different weeks (6 non-critical, 9 critical) to understand individual SARS-CoV-2 specific TCR repertoire evolution. For this subset, log2 fold change of TCR frequencies between consecutive data points was calculated (Fig.S5-a, b). This analysis revealed that the majority of non-critical (5/6) and critical (7/9) patients experienced decline in the frequency of CoV-common TCRs at least once during their active COVID-19. 33% (2/6) of non-critical and 56% (5/9) of critical patients didn’t have any changes in the depth of SC2-unique TCR repertoires. In all individuals, SC2-unique and CoV-common TCR repertoires were changing in the opposite directions 9 times, 5 times in the same direction and 8 times only one of the repertoires was changing between consecutive weeks while the other remained the same highlighting that TCR repertoires are constantly evolving. Noteworthy, the range of the magnitude of the change was comparable between SC2-unique and CoV-common TCR frequencies and between critical and non-critical groups. Facing the limitation of our dataset wherein only 1 or 2 data points had been collected for most patients (Fig.S1-c), we attempted to extrapolate general group trends from the obtained longitudinal and single-point observations. Despite intra- and interpersonal variability of SARS-CoV-2 TCR repertoire expansion and contraction (Fig.S6), there seemed to be some overall trends. In particular, multiple patients from both disease severity groups had a rise of SC2-unique and CoV-common TCR frequencies around weeks 2-3 (Fig.S7A,B). The depth of CoV-common TCR repertoires also seemed to increase around week 6 in some individuals with critical and non-critical COVID-19 severity (Fig.S7A,B). In the critical group, SC2-unique TCRs showed a tendency to increase in their frequency throughout the entire duration of the study despite sustained T-cell depletion (Fig.7B, Spearman rho=0.41, p=0.02). This was not the case for the non-critical patient group (Fig.7A, Spearman rho=-0.06, p=0.71) where the maximum frequency of SC2-unique TCRs was reached by weeks 2-3 (Fig.S7A).

**Figure 7.**
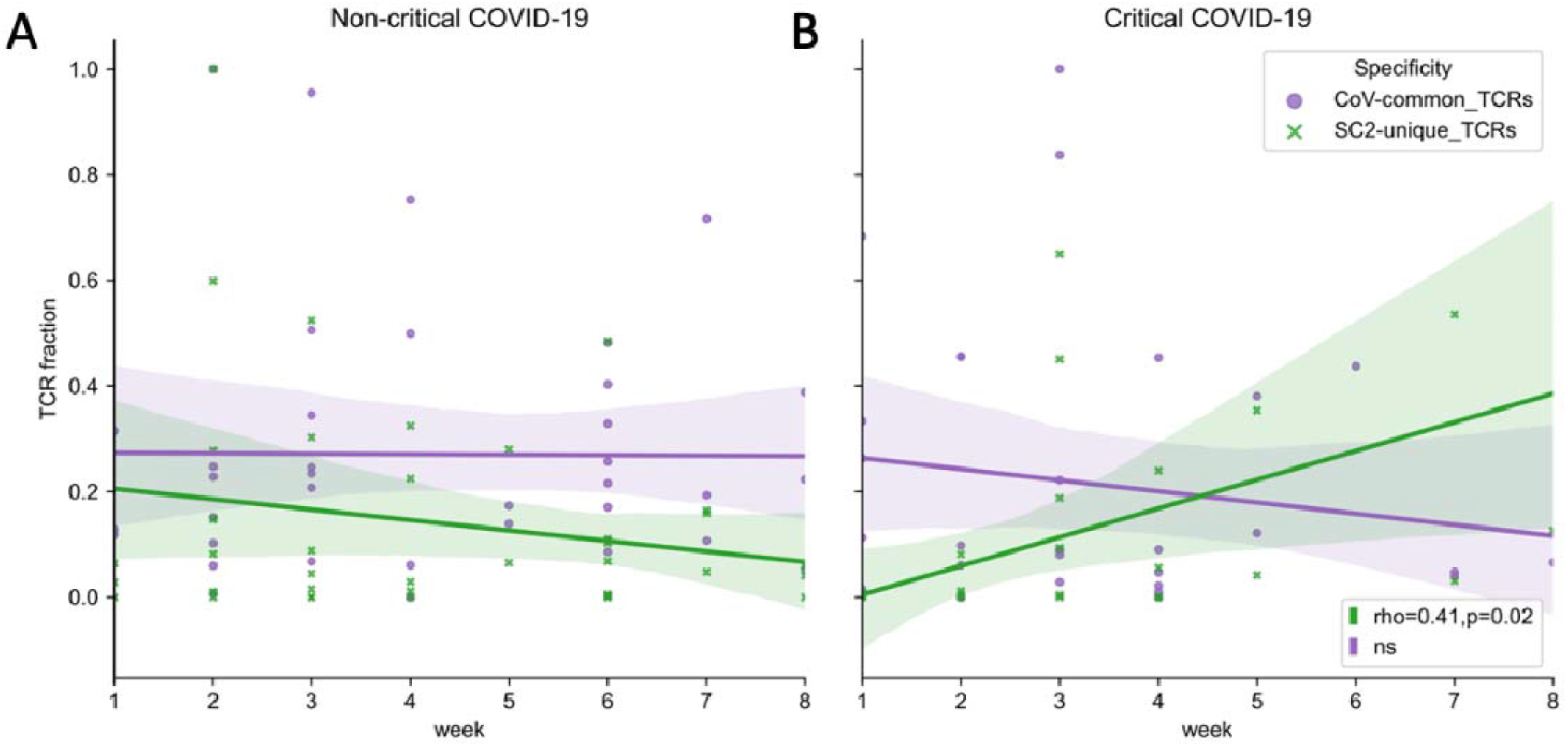
(**A**) Symptomatic non-critical patients demonstrated significant increases of neither SC2-unique (green, Spearman rho=-0.06, p=0.71) nor CoV-common (purple, Spearman rho=0.06, p=0.72) TCR repertoires when disease and recovery stages were evaluated together. (B) In contrast, relative frequencies (depth of the repertoire) of SC2-unique TCRs (green, Spearman rho=0.41, p=0.02) but not CoV-common TCRs (purple, Spearman rho=-0.10, p=0.58) were increasing significantly during disease progression in critical patients. Lines represent an estimate of the central tendency of multiple inter- and intra-individual values combined within each disease severity group (n = 14 critical, 32 non-critical) with a 95% confidence interval for those estimates shown as shadow areas in case multiple data points were available at overlapping time points.

The most prominent breadth changes occurred exclusively in SC2-unique TCR repertoires, while the breadth of TCR repertoires reactive to CoV-common epitopes remained relatively stable in both patient groups during the entire study period (Fig.S7C,D). Notably, the tendency for the increased breadth of SC2-unique TCRs was dominated by non-critical patients and mostly occurred within the first two weeks after symptom onset (Fig.S7C). This trend was delayed in the critical group and was supported by only few individuals (Fig.S7D).

Those findings reinforce that it is the build-up of the SARS-CoV-2 TCR repertoires, which is happening during the first two weeks of the disease, that might be crucial for differentiating COVID-19 severity. The SARS-CoV-2 specific CD8+ TCRs in critical patients seem to expand slower than in non-critical patients.

## DISCUSSION

Notwithstanding the general agreement on the importance of T cells during SARS-CoV-2 infection, the contribution of SARS-CoV-2-unique and cross-reactive T-cell responses towards modulation of COVID-19 severity remains not fully resolved. In this study, we combined our newly generated TCR sequences from COVID-19 patients hospitalized in a single center in Belgium with public datasets to gain insights into the specificity and evolution of CD8+ TCR repertoires in critical and non-critical symptomatic COVID-19 patients.

We observed that CD8+ T cells predicted to target CoV-common epitopes are mounted, despite T-cell depletion, in both critical and non-critical patients during the first week after symptom onset. Since the frequency of SC2-unique TCRs was significantly lower in our study samples at that time, we deduced that the depth of CoV-common TCR repertoires was higher in the first week due to more rapid clonal expansion of pre-existing cross-reactive memory CD8+ T cells. Thus, if an individual has encountered common cold coronaviruses before, memory CD8+ T cells are formed and may be the first to respond upon reactivation with CoV-common epitopes from SARS-CoV-2 while *de novo* induced SC2-unique CD8+ T-cell immunity has not been developed yet. This explanation falls in line with previous reports where T cells recognizing SARS-CoV and seasonal coronaviruses were found to be crossreactive to SARS-CoV-2 (Braun et al., 2020; Lineburg et al., 2021; Mateus et al., 2020; Nesterenko et al., 2021). Alternatively, individuals may have had previously developed crossreactive CoV-common CD4+ T cells that might have facilitated more rapid CD8+ T cell development at the beginning of the SARS-CoV-2 infection, as has been recently demonstrated to be the case for antibody response to vaccination (Elias et al., 2022).

By the end of the second week after symptom onset, activation and expansion of SC2-unique TCRs seem to have occurred in non-critical patients despite T-cell depletion. This conclusion can be made since at week 2 but not week 1 we observed (1) a significant difference in the frequency of those TCRs between critical and non-critical patients; (2) no significant difference in the depth of SC2-unique and CoV-common repertoires and (3) increase in the median number of putatively recognized SC2-unique and CoV-common epitopes in non-critical patients. In contrast, critically ill patients experienced reduction in both the breadth of CoV-common and SC2-unique TCR repertoires and respective response diversities at this time. This disparity can partially be explained by lymphopenia which is known to be more pronounced in critical patients (Tan et al., 2020) and seems to affect CD8+ T-cell population more (Chen and John Wherry, 2020; Liu et al., 2020). Previous research has provided evidence that T cells are dying during severe COVID-19 due to apoptosis (André et al., 2022), but the specificity of those T cells has not been addressed yet. We further speculate that during lymphopenia in critical patients, SARS-CoV-2 recognizing rather than any CD8+ TCRs might be specifically depleted. If random TCRs were dying, the breadth of SARS-CoV-2 TCR repertoires would not have decreased as specific TCRs constitute the minority of all unique TCRs in the repertoire. When longitudinal data were analyzed (8 weeks), a significant increase in the depth of SC2-unique TCR repertoires was also detected in critically ill patients, which is especially interesting in light of persisting T-cell depletion.

Overall, our findings align with previously proposed mechanisms of effective T-cell response development, where timely (within two weeks) activation and expansion of T cells allows control of the virus, corresponding to a milder form of COVID-19 (Moss, 2022; Sette and Crotty, 2021). Conversely, if the activation and/or expansion of the SARS-CoV-2 specific T-cells is delayed, the virus multiplies unchecked, and the overactivated immune system results in inflammation worsening the symptoms (Tan et al., 2021). Furthermore, we have observed the dominance of CD8+ T cells putatively recognizing SC2-unique rather CoV-common epitopes, which supports the previous report that in contrast to CD4+ T cells, most expanded CD8+ T cells did not cross-react with seasonal coronaviruses (Ferretti et al., 2020). Additionally, considering slight underrepresentation of SC2-unique epitopes in comparison to CoV-common in the trained TCRex models (19 SC2-unique vs 28 CoV-common), the dominance of SC2-unique TCRs in the evolution of T-cell response hints towards the relative immune response contribution of newly expanding T-cell clones compared to potentially preexisting cross-reactive T cell clones in COVID-19 patients.

Despite the congruence of our results, they should be interpreted with caution: only 4 out of 8 critical and 2 out of 8 non-critical patients had the data available for both the first- and second-week post symptom onset, and among those, not everyone followed described group trends. For instance, half of the critical patients with continuous data (2 out of 4) demonstrated elevated CoV-common TCR frequency at week 2, which hints that even patients with the same disease severity do not have a uniform T-cell immune response to SARS-CoV-2. For example, while pre-existing cross-reactive CD8+ T-cells may provide clinical protection to some individuals (Mallajosyula et al., 2021; Schulien et al., 2021), this protection has also been questioned (Chen and John Wherry, 2020; Saggau et al., 2022; Sette and Crotty, 2020). In agreement with the latter outlook (detrimental role of T cells), we have observed that disease outcome was fatal in three critical patients who had reached the highest frequencies of CoV-common TCRs during the first two weeks of COVID-19.

The observed interpersonal variation in the CD8+ T-cell response within the same group in our dataset could also be attributed to the fact that current TCRex models do not cover the entire epitope space available to CD8+ T cells. It has been experimentally evaluated that every individual recognizes 17 MHC-I SARS-CoV-2 epitopes (Tarke et al., 2021), which is more than was predicted with our recognition models. Hence, the magnitude of CD8+ T-cell response is likely to be underestimated in our analysis. Therefore, the response could become more uniform across patients once there is enough experimental data for more epitopes, and recognition models are built for them.

Another limitation inherent to the analysis of TCR repertoires from blood samples is that the migration of T cells from the circulation to tissues remains hidden. Consequently, the observed intragroup dissimilarities in SC2-unique TCR levels could be reflective of the differences in the distribution rather than in the speed of generation of the respective T cells. If the same number of SC2-unique T cells is generated equally promptly in both patient groups, but more of them are attracted to and retained at the site of the infection (lungs) in critical patients, our analysis cannot detect that. Further examination of TCR repertoires from recovered critical individuals could help shed light on this puzzle and reflect their potential to develop SC2-unique response more precisely.

Finally, it has been demonstrated that in case of some HLA-alleles, they strongly shape the response, resulting in a particular set of immunodominant T-cell epitopes (Francis JM et al., 2021; Lineburg et al., 2021; Wu et al., 2022). Thus, our models will not suffice to track the response unless the models for those epitopes are available. However, there is evidence that multi-epitope T-cell response mitigates the effect of viral escape mutations (Redd et al., 2021). The broad and redundant response is more important for SARS-CoV-2 control than the recognition of particular epitopes in our observations as well. Thus, monitoring the TCR repertoire depth and breadth and response redundancy and diversity in infected, vaccinated and recovered individuals could be helpful to better assess current or future protection and the necessity for an additional intervention to prevent critical disease or preserve the protection against reinfection and emerging variants. *In silico* approach can be leveraged to extract information such as specificity faster and with more flexibility than *in vitro* testing. Accordingly, our study offers a generalizable computational framework that complements the current standard of antibody-based testing of COVID-19 immunity with TCR repertoire analysis.

## Author contributions

P.M., K.V., K.L., B.O. and A.P. conceived and planned the experiments. A.V., T.d.B., L.v.P., M.v.F., I.B., E.B., C.V.D., C.T., S.H.v.E., E.V., E.B. collected samples and generated the experimental data. A.P. and P.M. developed the analytical framework and analyzed the data. A.P., W.A., G.V., B.O., K.L., K.V., and P.M. contributed to the interpretation of the results. A.P. took the lead in writing the manuscript. All authors reviewed the results, provided critical feedback and approved the final version of the manuscript.

## Acknowledgments

We thank Marianne Mangelschots, Sergio Garcia, Charlotte Drieghe, Cindy Van Hoyweghen and Niels Lonneville for excellent technical assistance.

## Data Availability Statement

The publicly available raw TCR sequencing data analyzed in this study were downloaded from iReceptor Gateway [https://gateway.ireceptor.org/login]. The processed dataset of CD4+/CD8+ COVID-19 TCR repertoires and epitope specificity predictions generated for this study can be found in the GitHub repository [https://github.com/apostovskaya/CovidTCRs/data].

## SUPPLEMENTARY MATERIALS

### 1 Supplementary Tables

Supplementary Material should be uploaded separately on submission. Please include any supplementary data, figures and/or tables.

**Table S1.** Summary of WHO criteria: COVID-19 disease severity in adults.

|  |  |  |
| --- | --- | --- |
| Mild | Symptomatic COVID-19 with: | No clinical evidence of viral pneumonia* or hypoxia. |
| Moderate | Clinical evidence of pneumonia* with: | RR ≤ 30 breaths/minute<br>AND<br>SpO<sub>2</sub> ≥ 90% on room air |
| Severe |  | RR > 30 breaths/minute<br>OR<br>SpO<sub>2</sub> < 90% on room air<br>OR<br>clinical signs of severe respiratory distress |
| Critical | ARDS with: | PaO<sub>2</sub>/FiO<sub>2</sub> ≤ 300 mmHg<br>With PEEP or CPAP ≥ 5 cmH<sub>2</sub>O<br>OR<br>need for invasive ventilation |

\*Clinical evidence of pneumonia: fever, cough, dyspnoea, fast breathing
Abbreviations: SpO<sub>2</sub>: oxygen saturation, PaO<sub>2</sub>/FiO<sub>2</sub>: arterial oxygen partial pressure (PaO<sub>2</sub> in mmHg) to fractional inspired oxygen (FiO<sub>2</sub>), RR: Respiratory rate, ARDS: acute
respiratory distress syndrome, PEEP: Positive end-expiratory pressure, CPAP: Continuous Positive Airway Pressure

**Table S2.**
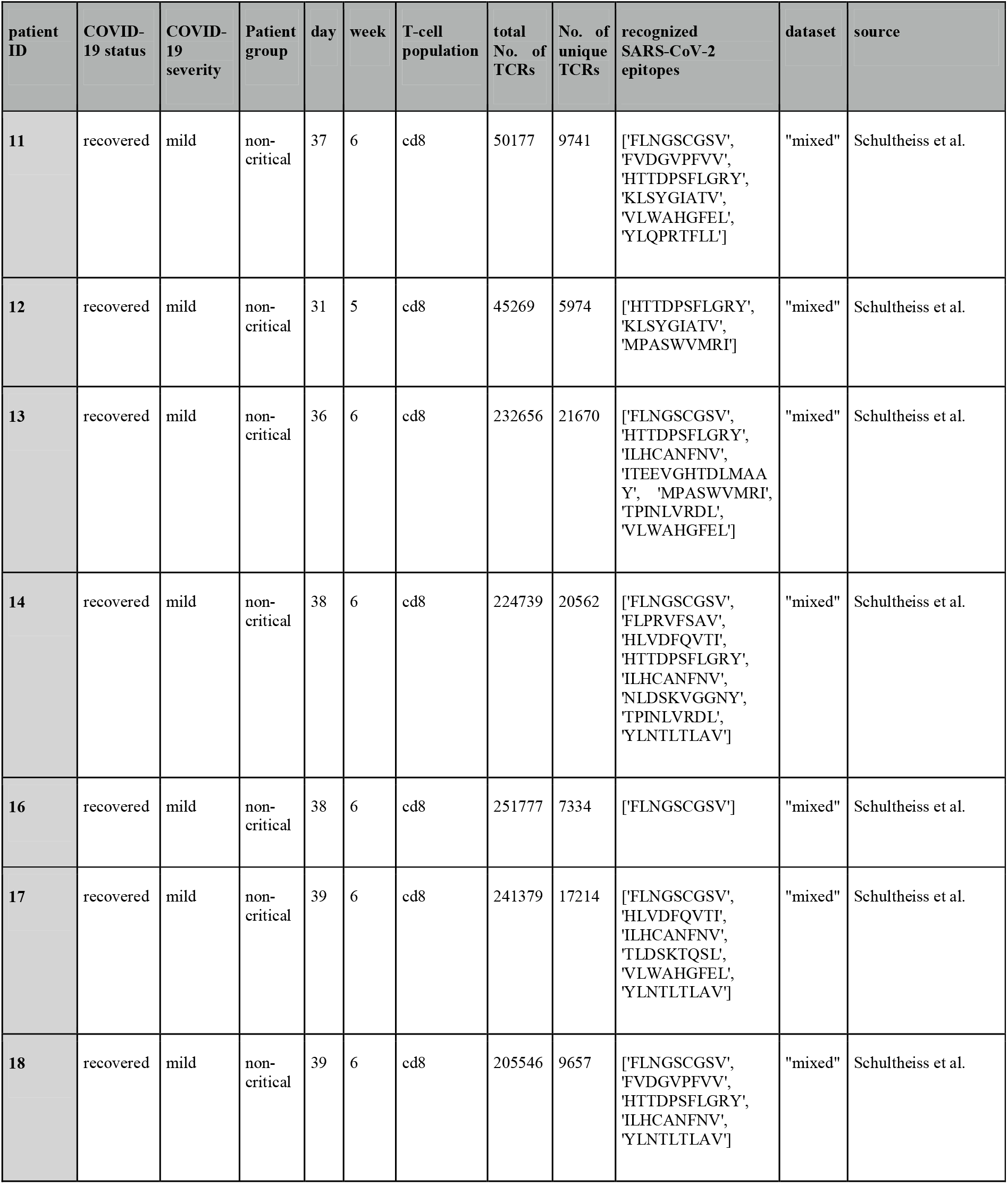
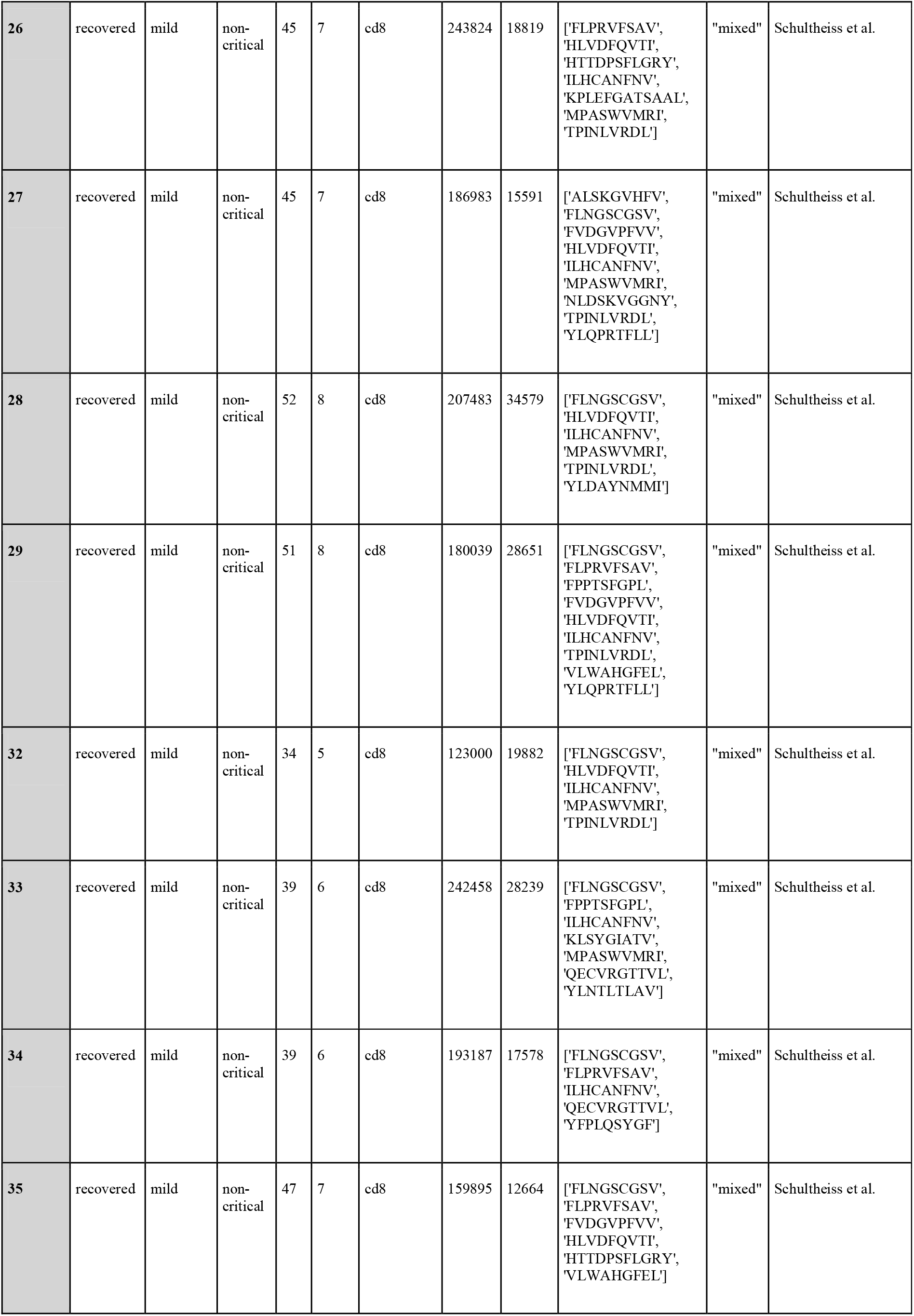
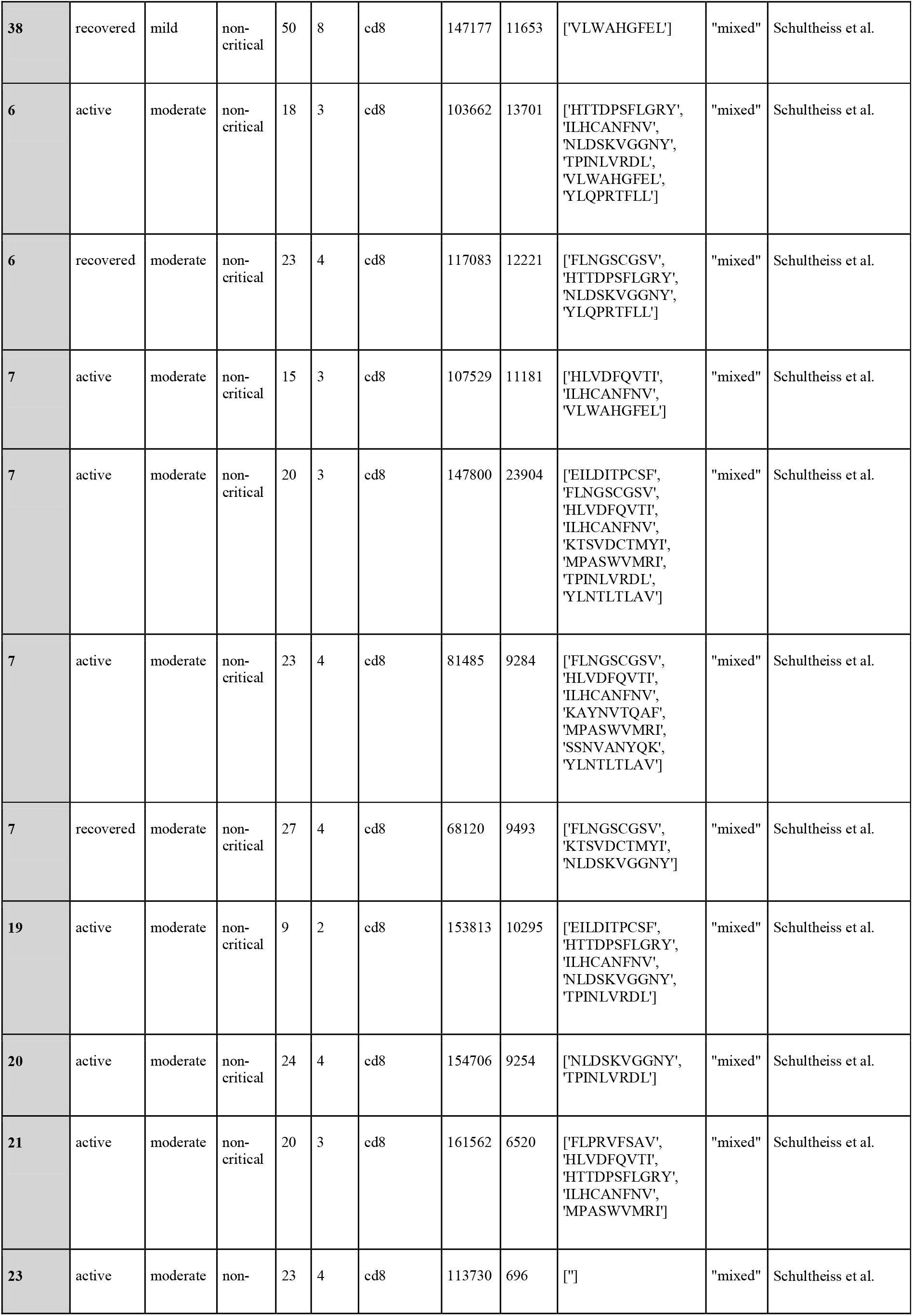
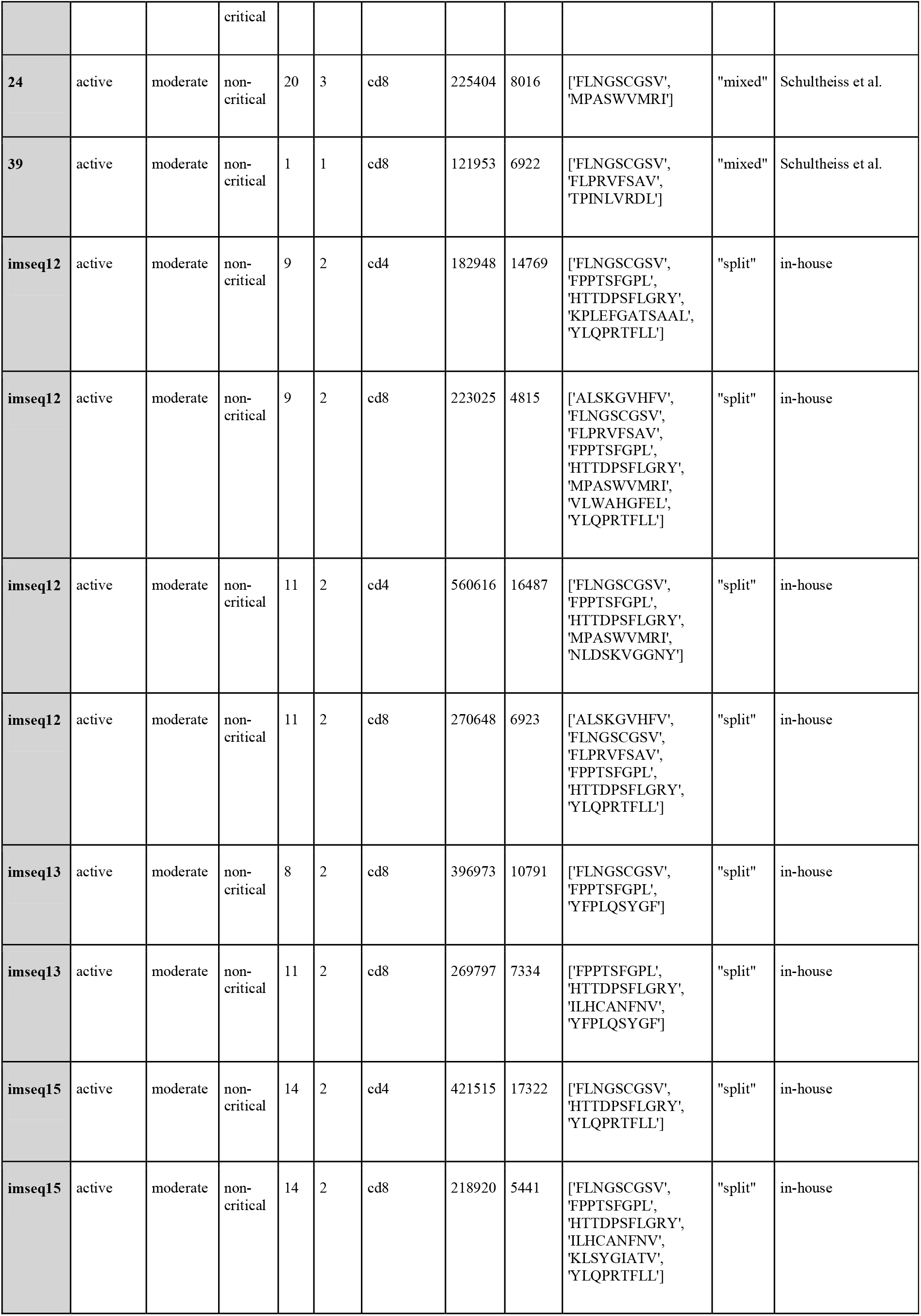
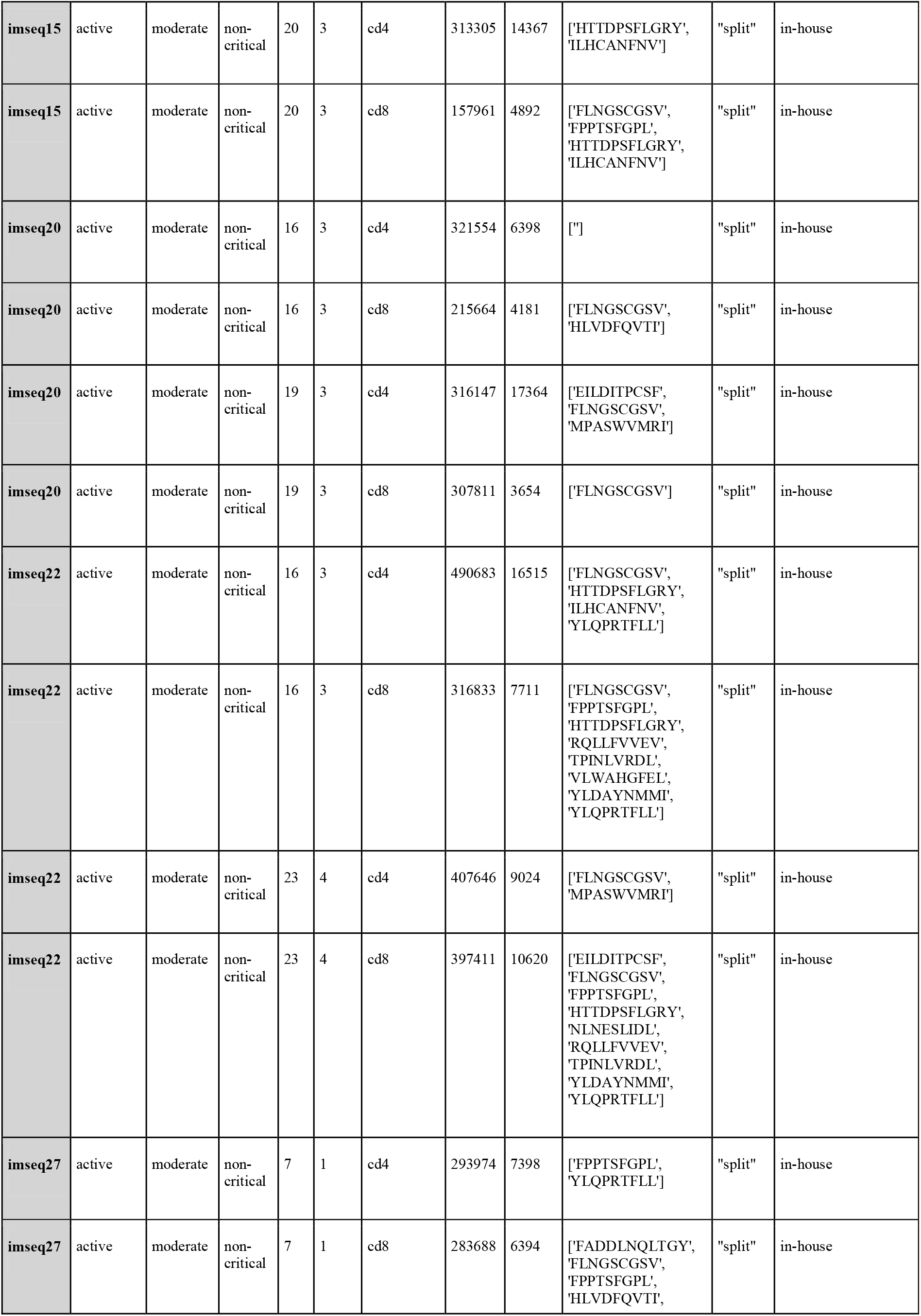
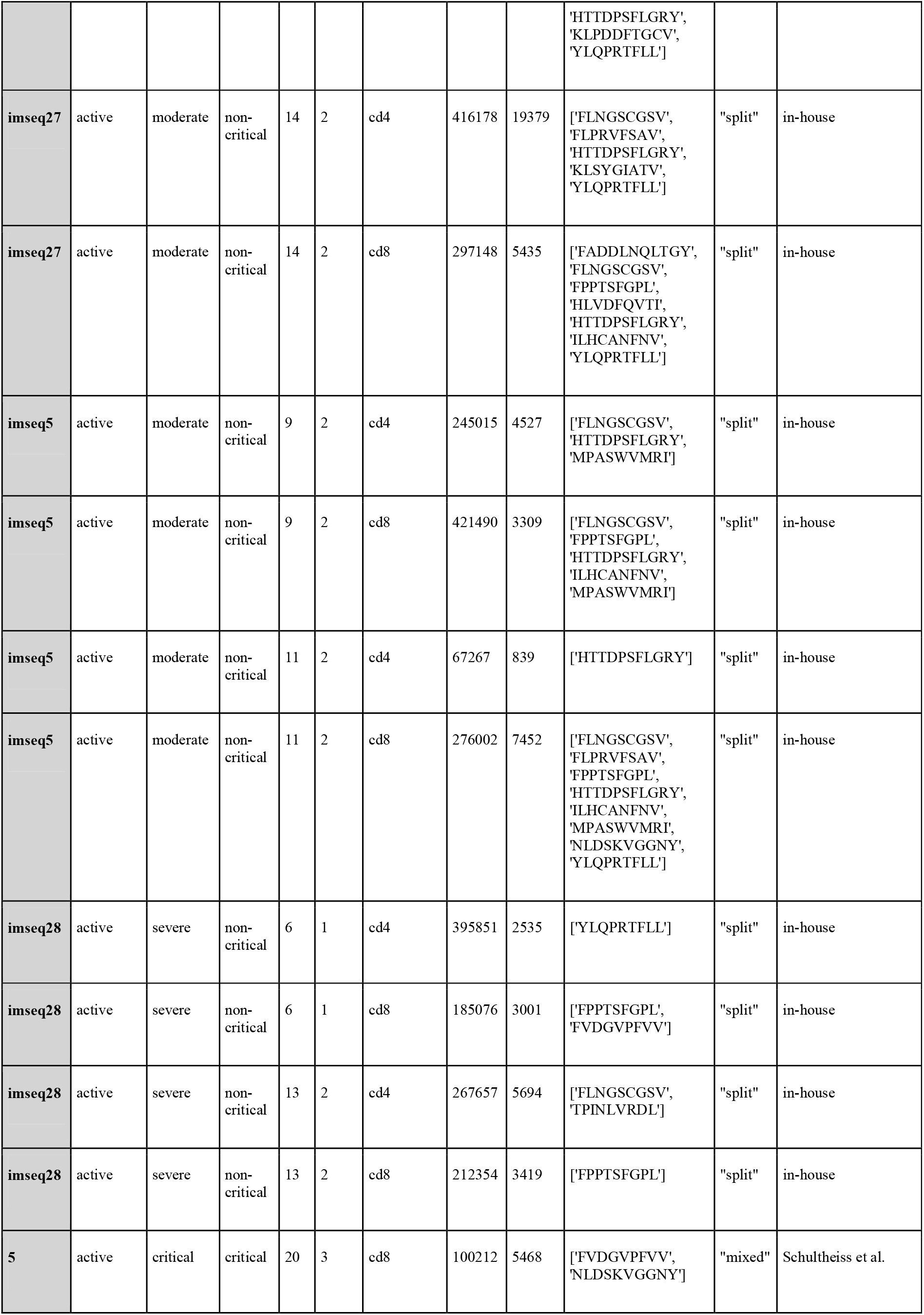
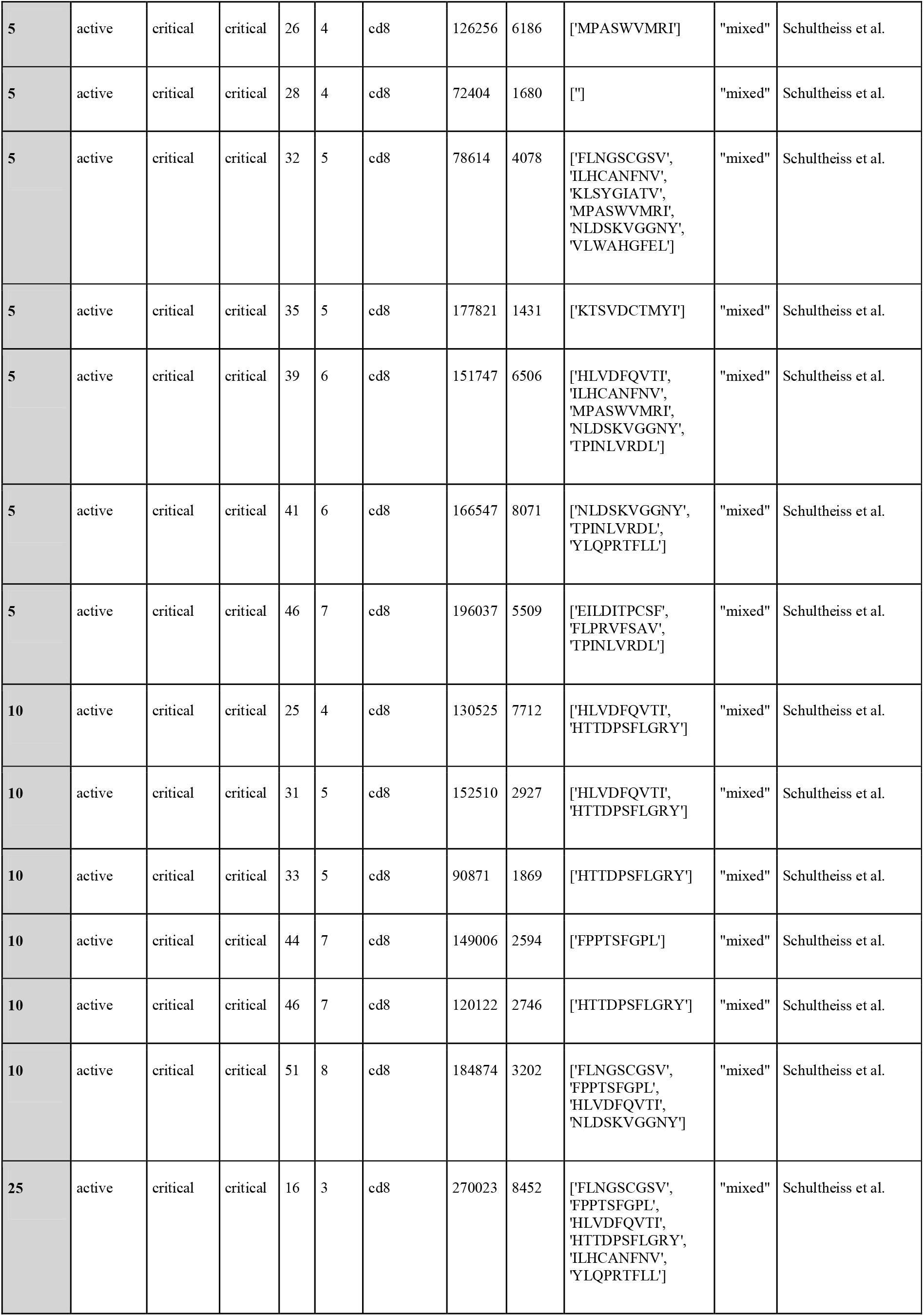
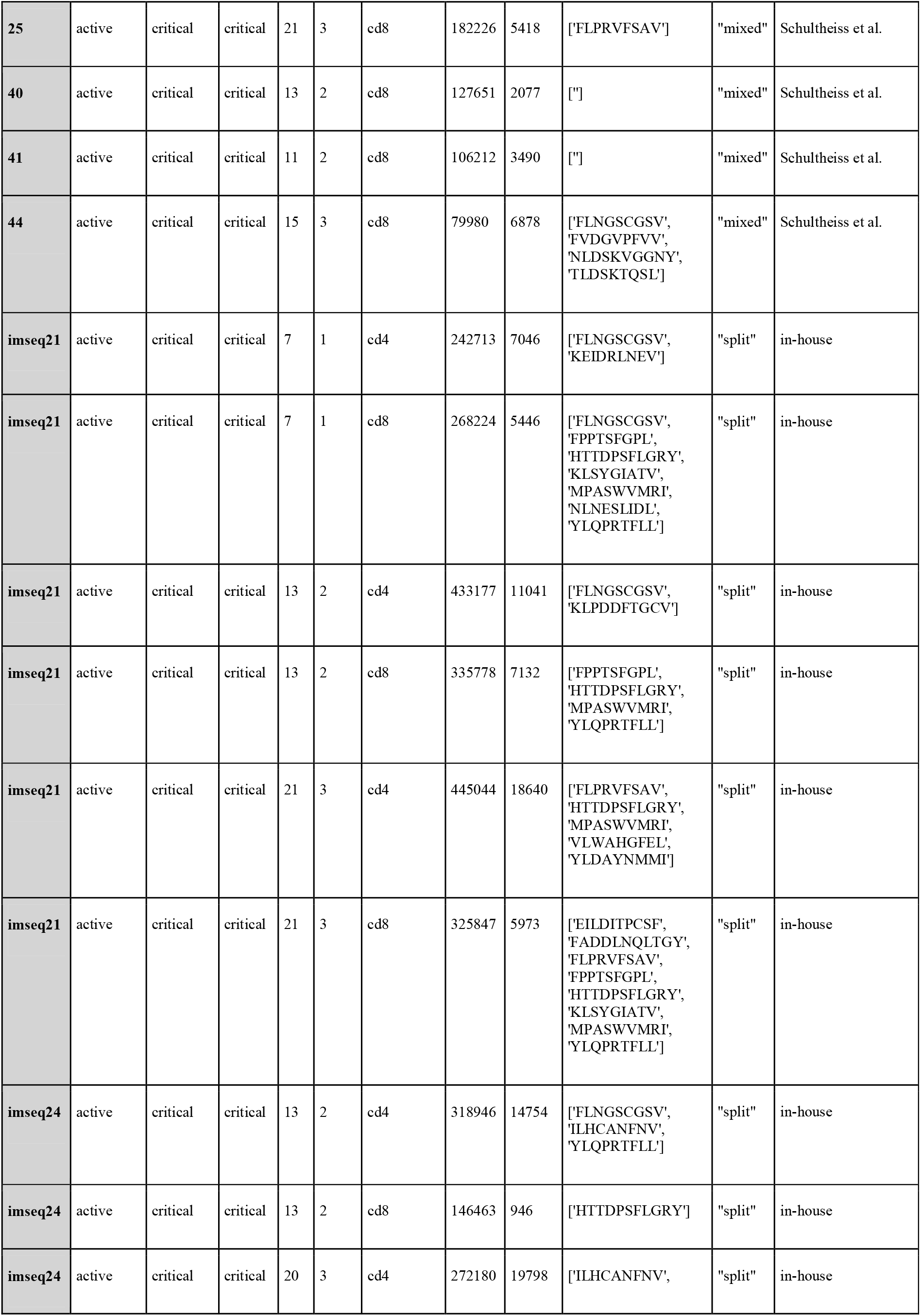
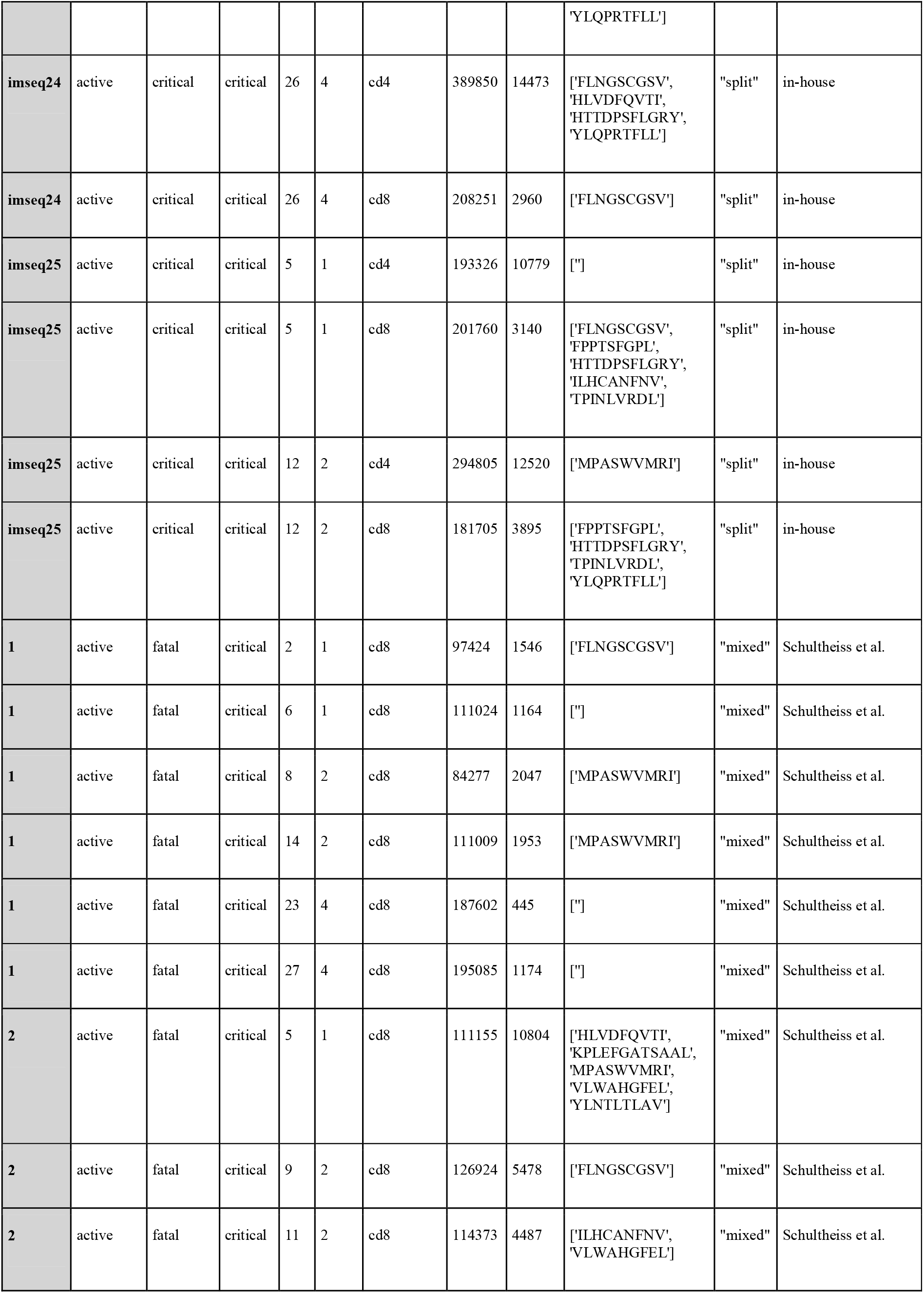
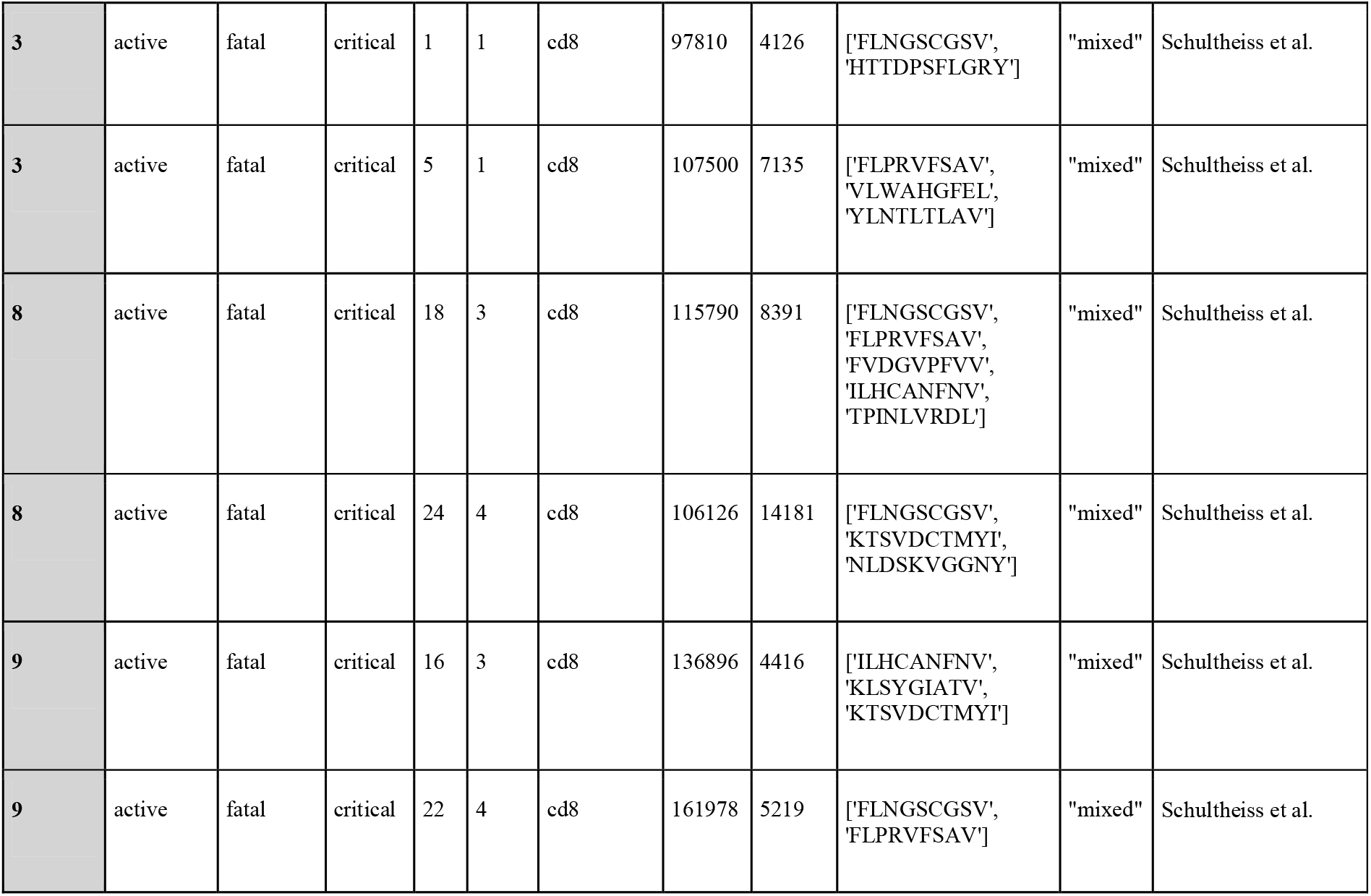
Patient and sample characteristics. The data is ordered by the COVID-19 severity and grouped by a patient. Time (day and week) refers to the time after the start of symptoms.

**Table S3.**
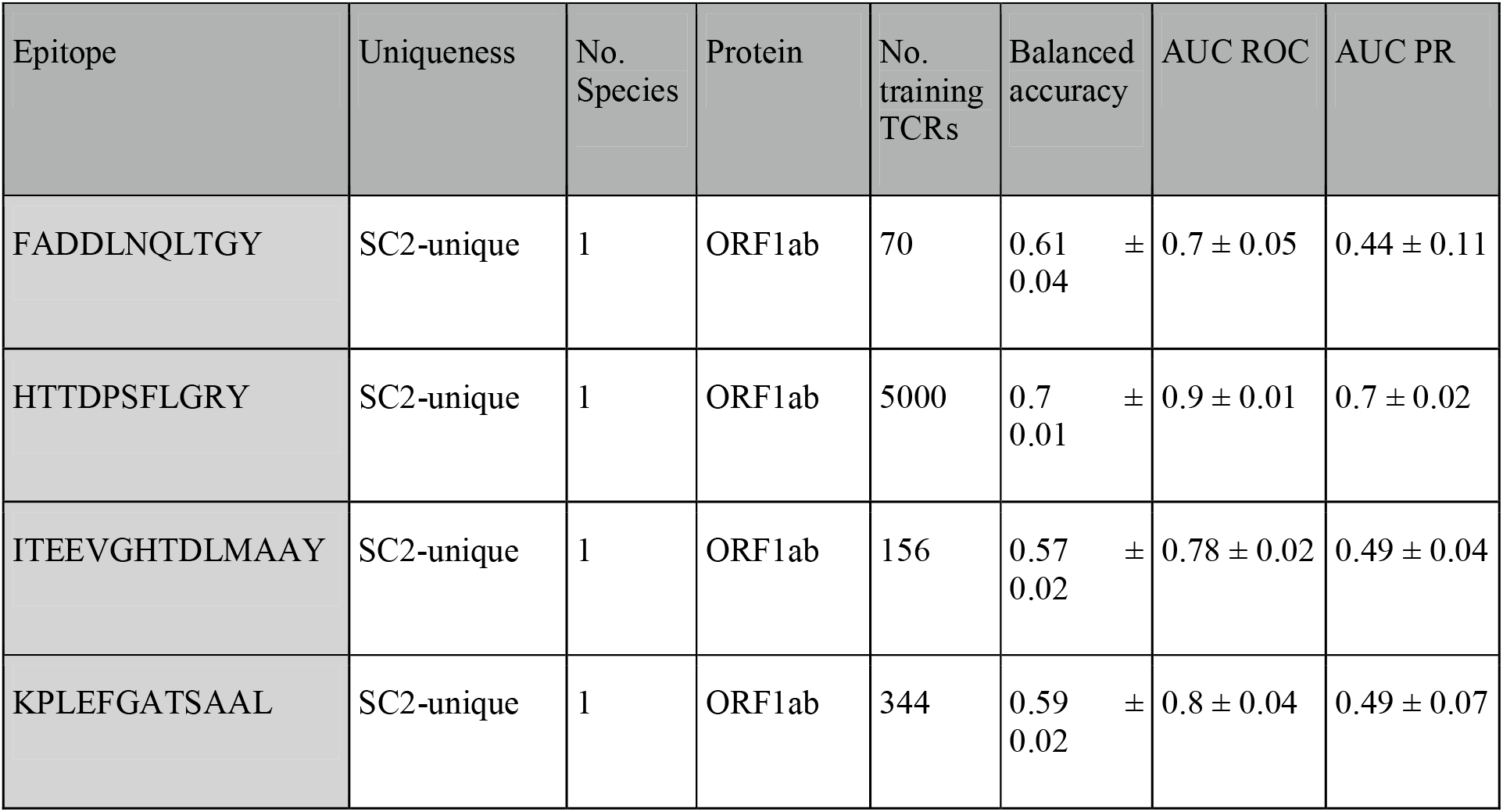
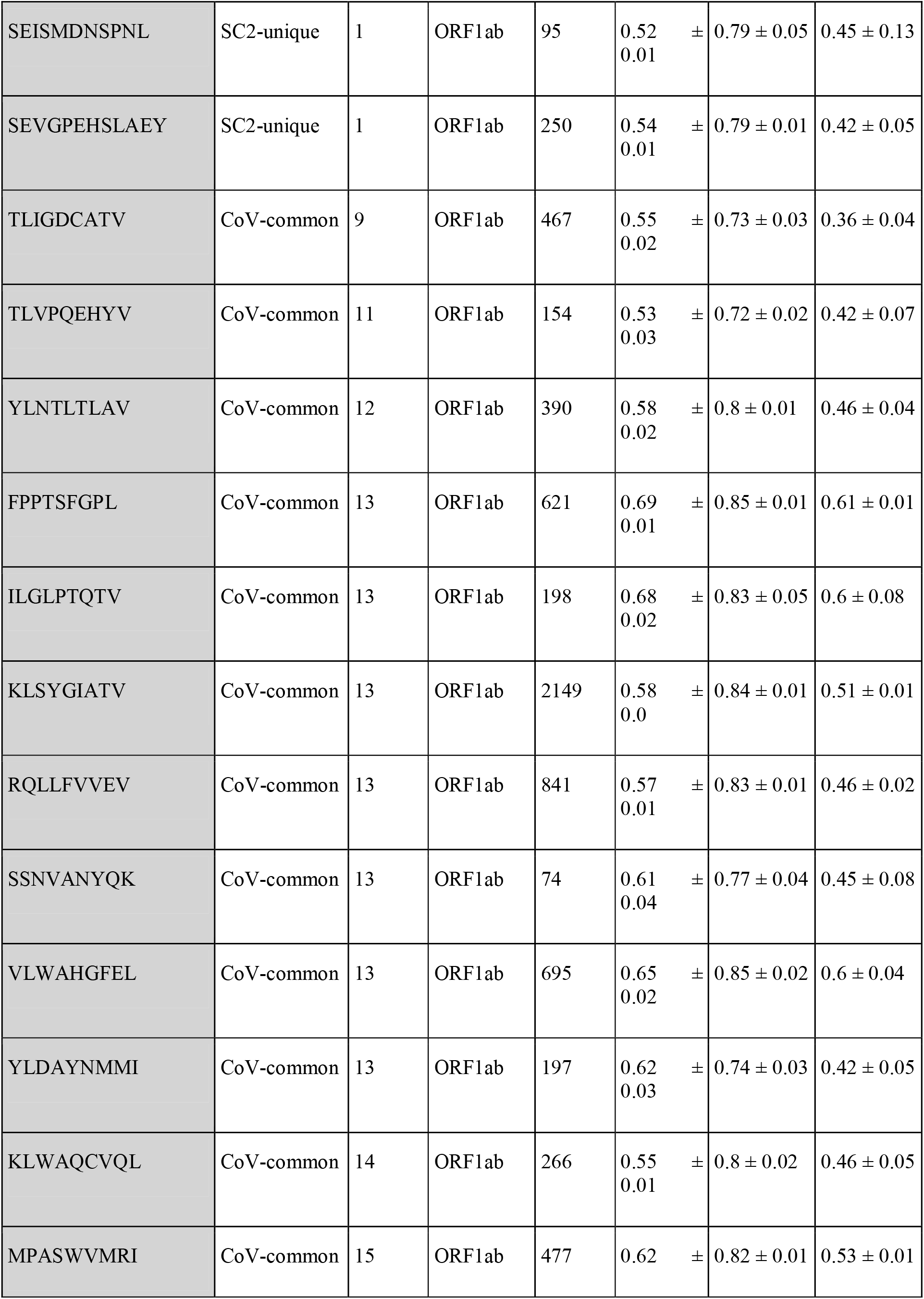
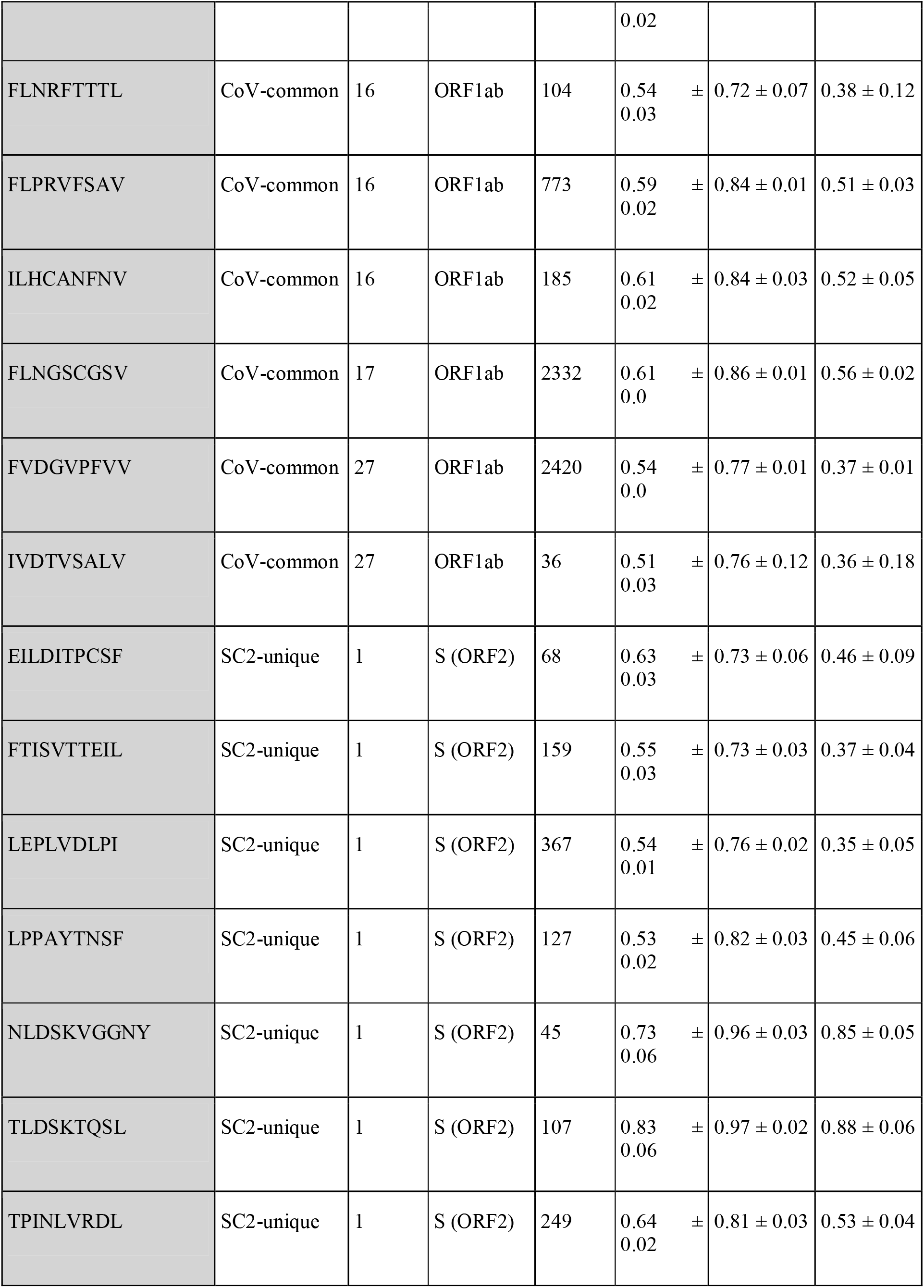
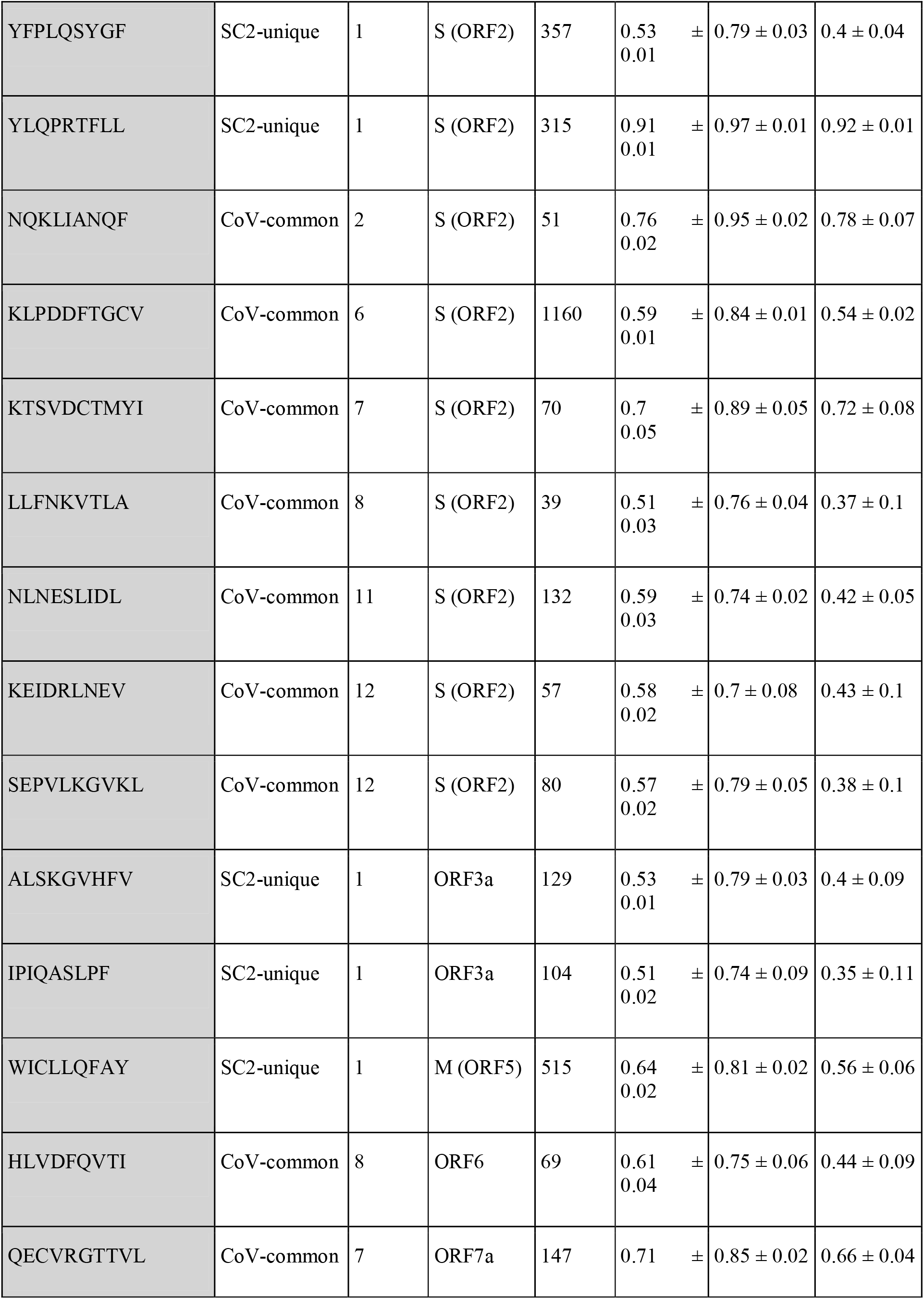
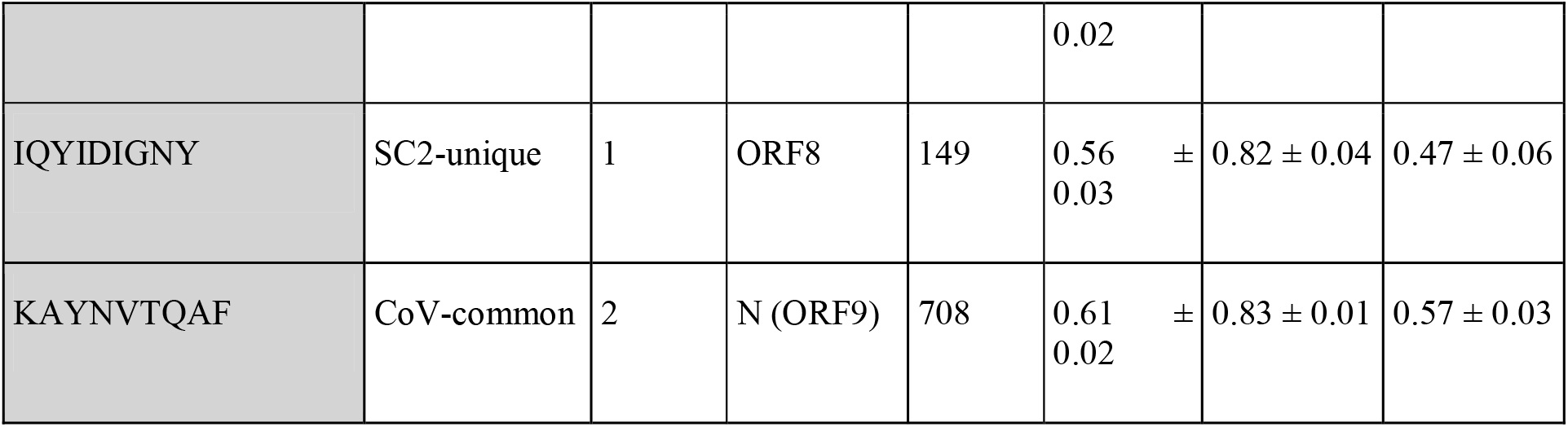
SARS-CoV-2 epitope recognition model statistics. The data is grouped by protein and ordered by epitope uniqueness, i.e., in how many species, including SARS-CoV-2, an epitope is present (No. Species).

**Table S4.** Results of Mann-Whitney U test between the depth of SC2-unique and CoV-common TCR repertoires of critical and non-critical active patients at different weeks. P-values are corrected for multiple comparisons (Bonferroni). Significant p-values are in bold (alpha is 0.05).

| week | adj_p-value critical | AUC critical | No. critical | adj_p-value non-critical | AUC non-critical | No. non-critical |
| --- | --- | --- | --- | --- | --- | --- |
| 1 | **0.049** | 1 | 5 | 0.162 | 1 | 3 |
| 2 | 1 | 0.551 | 7 | 1 | 0.49 | 7 |
| 3 | 1 | 0.611 | 6 | 0.193 | 0.776 | 7 |
| 4 | 1 | 0.639 | 6 | 1 | 0.52 | 5 |
| 5 | 1 | 0.75 | 2 | NA | NA | NA |
| 6 | 1 | 0 | 1 | NA | NA | NA |
| 7 | 1 | 0.5 | 2 | NA | NA | NA |
| 8 | 1 | 0 | 1 | NA | NA | NA |

**Table S5.** Results of Mann-Whitney U test between critical and non-critical patients. P-values are corrected for multiple comparisons (Bonferroni). Significant p-values are in bold (alpha is 0.05).

| week | parameter | adj_p-value | AUC |
| --- | --- | --- | --- |
| **1** | total_TCRcount | 0.742 | 0.733 |
|  | %unique_TCRs | 1 | 0.533 |
|  | Freq_CoV-common_TCRs | 1 | 0.467 |
|  | Freq_SC2-unique_TCRs | 0.907 | 0.667 |
| **2** | N_CoV-common_Eps | **0.026** | 0.796 |
|  | normN_CoV-common_Eps | 0.054 | 0.816 |
|  | N_SC2-unique_Eps | 0.054 | 0.714 |
|  | normN_SC2-unique_Eps | 0.433 | 0.592 |

### 2 Supplementary Figures

**Supplementary Figure 1.**
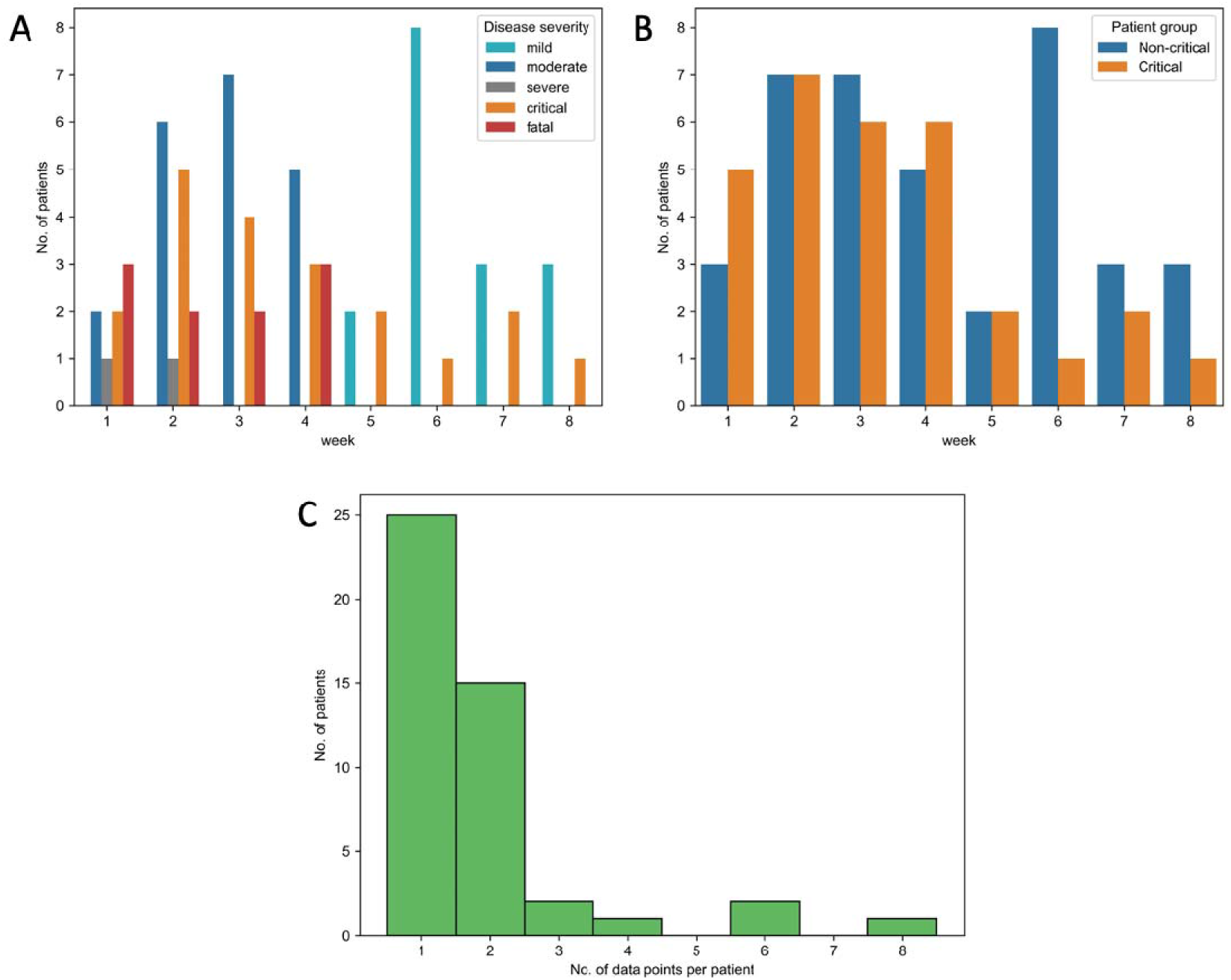
The distribution of the available data points in the analyzed patient cohort (merged dataset) between individuals and weeks of the study. (a) The number of available patients at every week of the study had a high variation between different disease severity groups. (b) Once patients were divided into critical (critical, fatal) and non-critical (mild, moderate, severe) groups based on their disease severity, the number of available patients became comparable between those two groups at most weeks of the study. (c) Most of the patients in the merged dataset had 1-2 available data points.

**Supplementary Figure 2.**
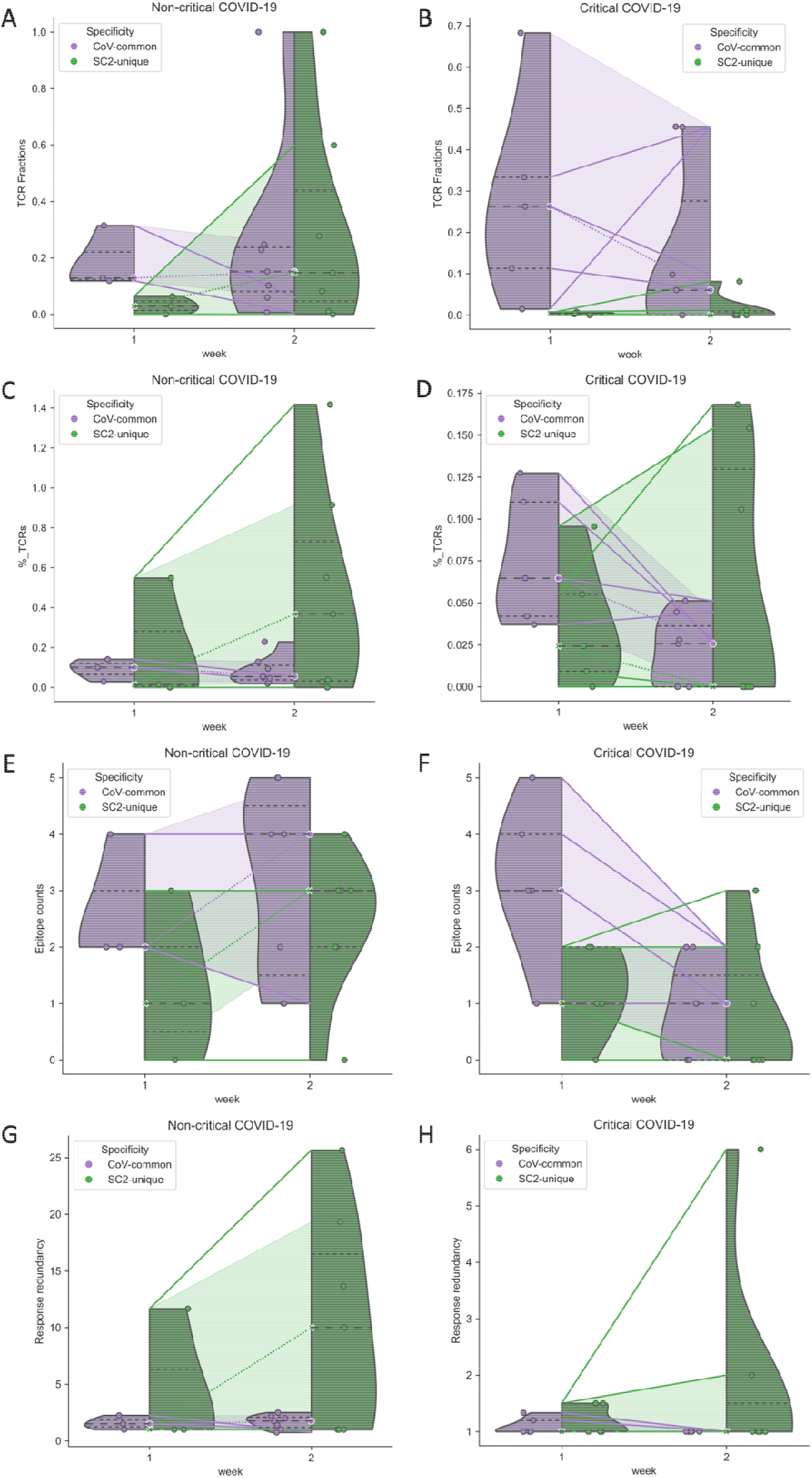
The changes in the SC2-unique and CoV-common CD8+ TCR repertoires of critical and non-critical patients that occurred between the first and the second weeks of COVID-19: repertoire depth (a, b); repertoire breadth (c, d); response diversity (e, f). Dotted lines indicate a change in the median values (median data points are encircled with white); solid lines correspond to the dynamics of individual patients.

**Supplementary Figure 3.**
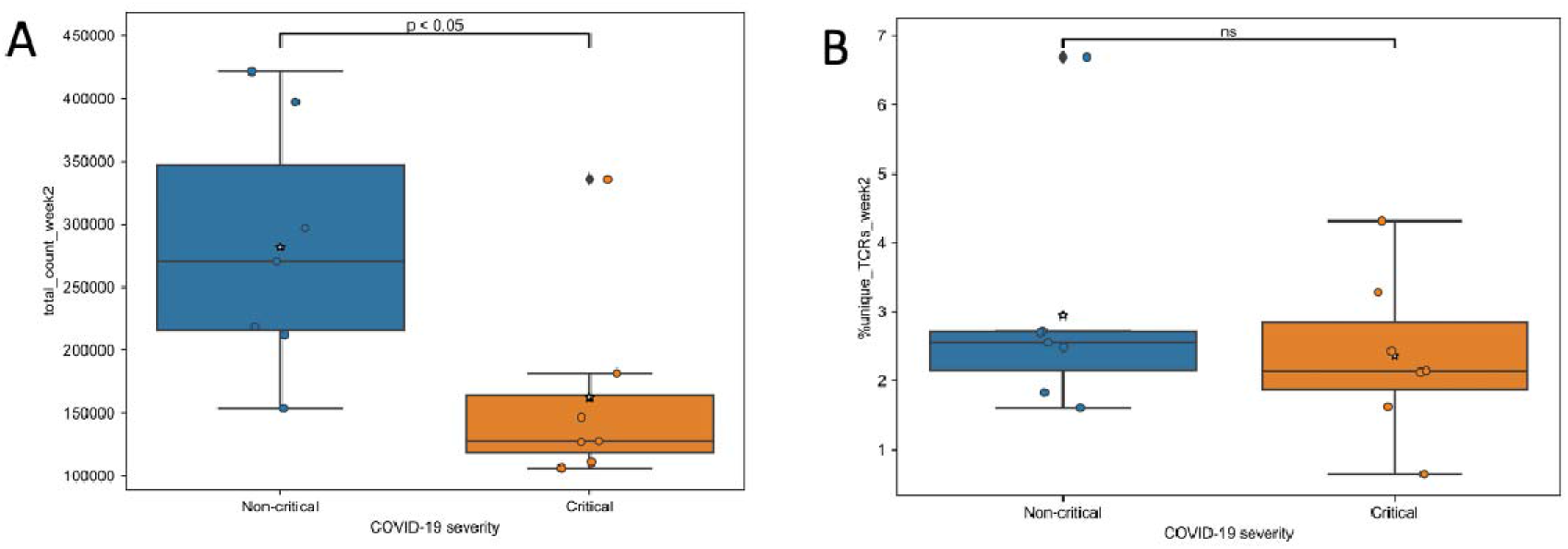
During the second week of COVID-19, the total number of TCRs (a, Bonferroni corrected Mann–Whitney U test p=0.033) but not the percent of unique TCRs (b, Bonferroni corrected Mann–Whitney U test p=0.783) were significantly higher in non-critical compared to critical patients. Mean values are represented by a star.

**Supplementary Figure 4.**
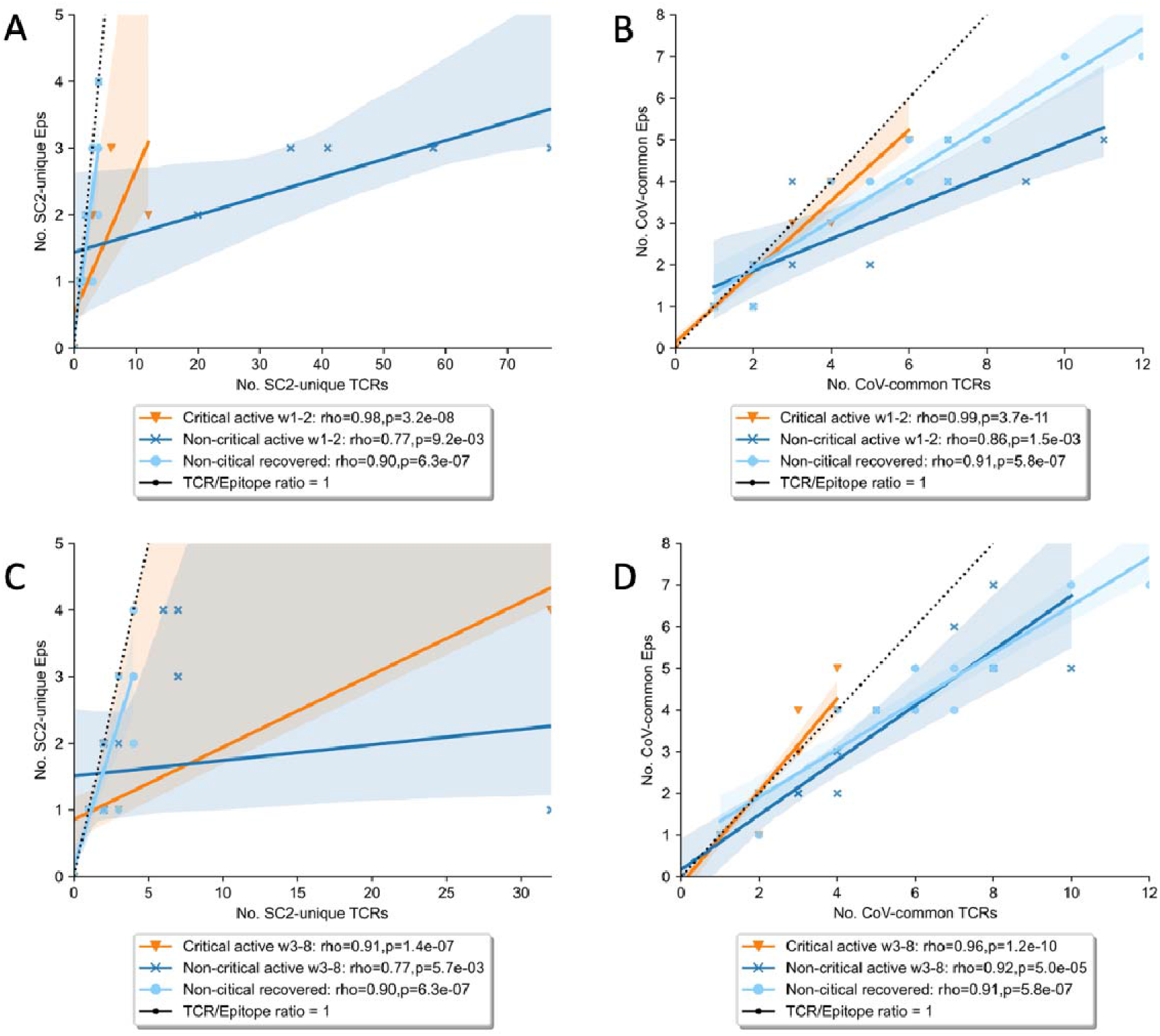
The diversity and redundancy of the response during the initial stages of the disease (a, b: weeks 1-2 after symptom onset), late stages of the disease and recovery stage (c, d: weeks 3+ after symptom onset) differed between patients with critical and non-critical COVID-19.

**Supplementary Figure 5.**
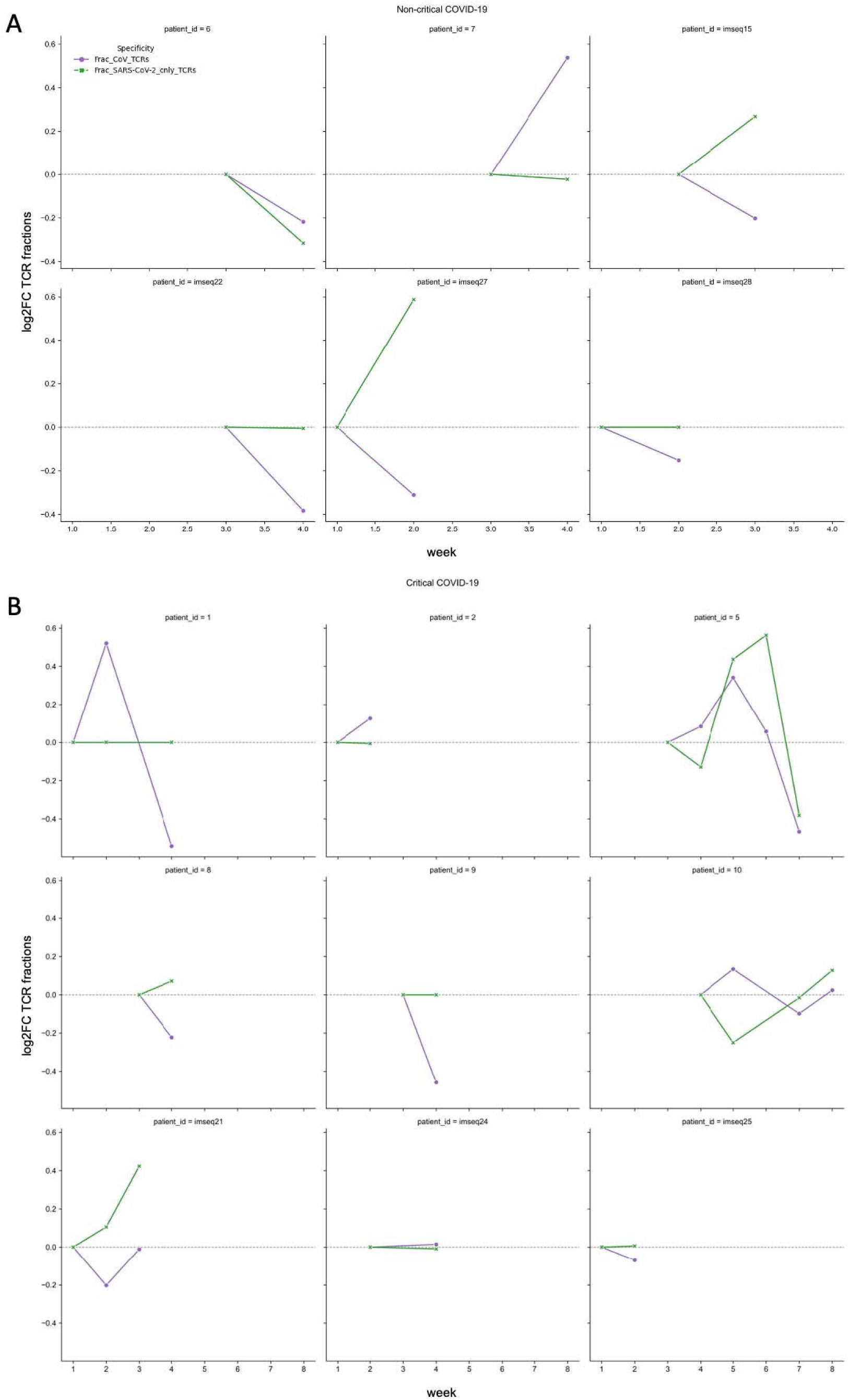
The magnitude of the alterations in putative CoV-common (purple) and SC2-unique (green) TCR repertoires, expressed as log2 fold changes in the relative frequencies of TCRs (depth of the repertoire) in non-critical (a) and critical (b) COVID-19 patients. Data points of the same patient are connected with lines to visualize temporal dynamics.

**Supplementary Figure 6.**
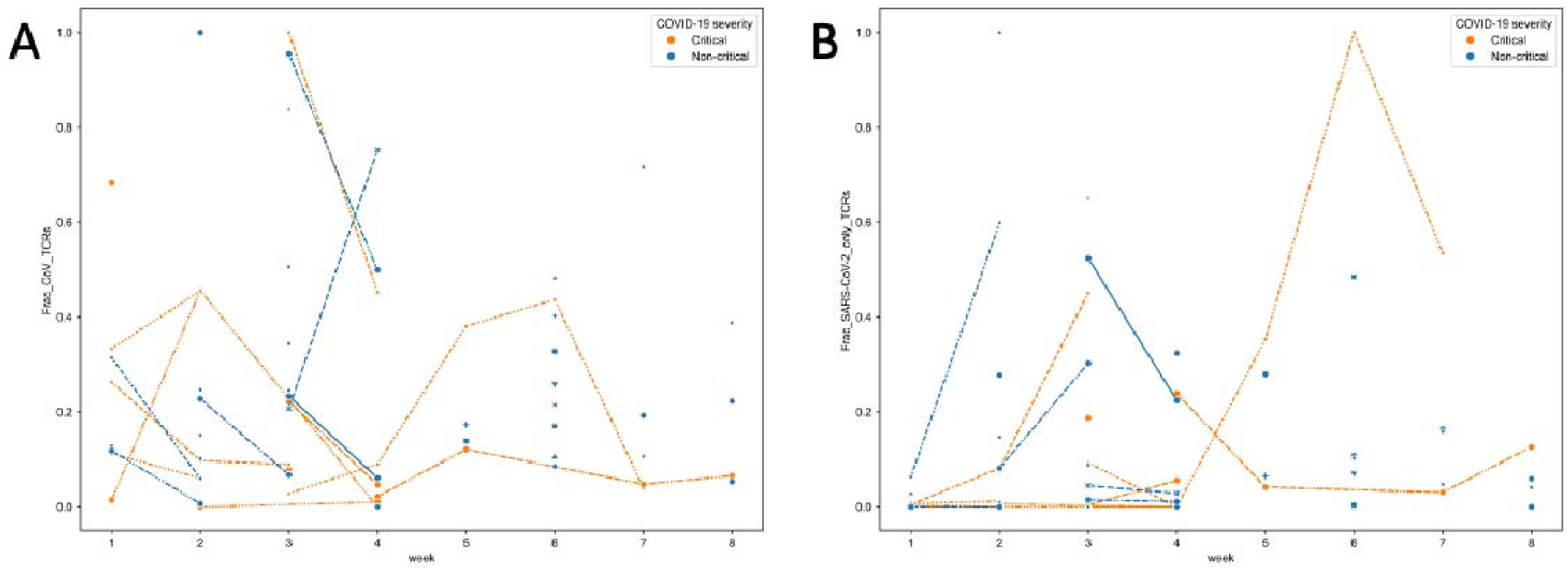
The evolution of putative CoV-common (a) and SC2-unique (b) TCR repertoires over time in individual critical (orange) and non-critical (blue) patients, expressed as changes in the relative frequencies of TCRs (depth of the repertoire). Raw data points of the same patient are connected with lines to visualize longitudinal dynamics.

**Supplementary Figure 7.**
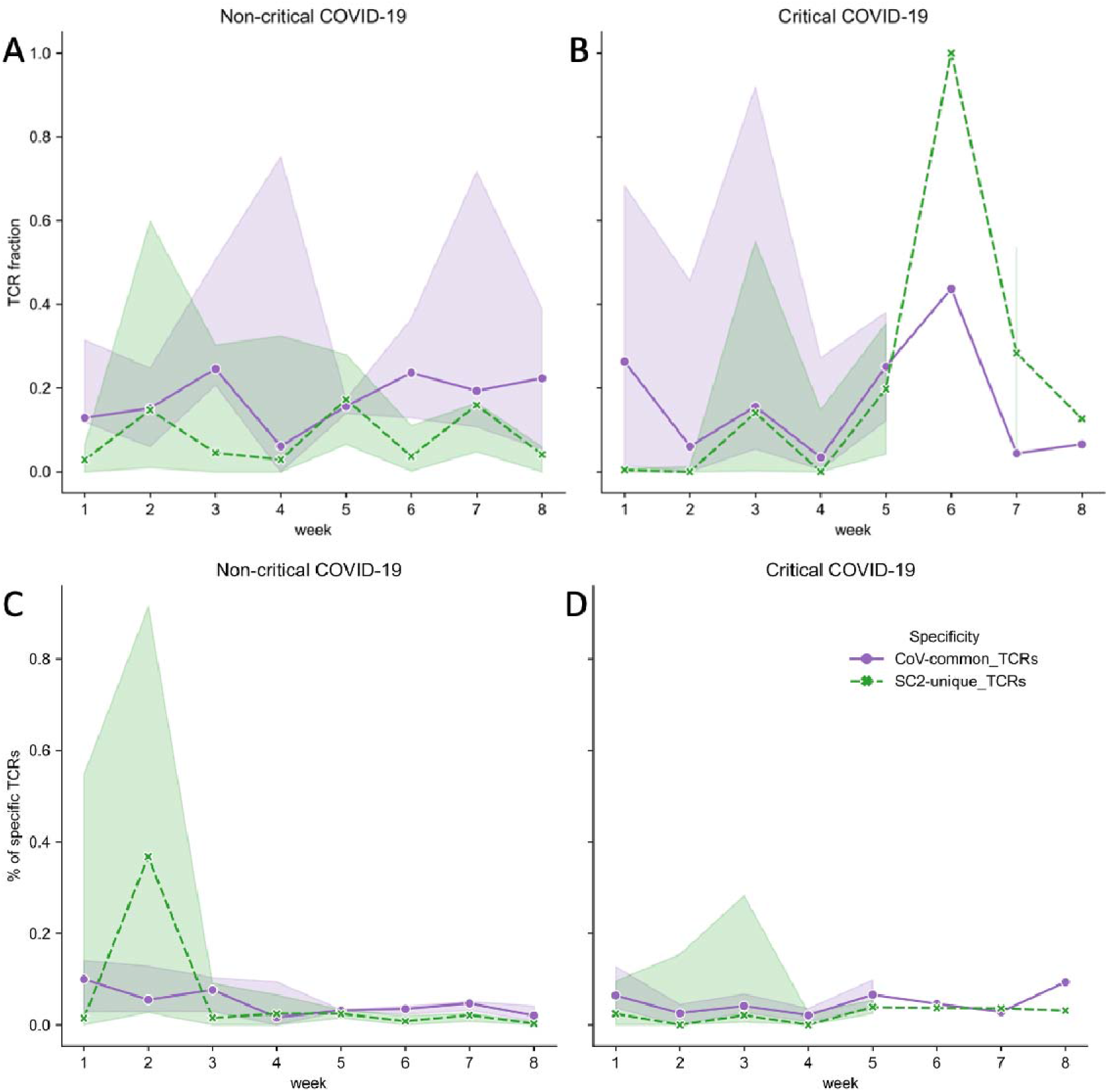
The dynamics of putative SARS-CoV-2 T cells reactive to epitopes unique to SARS-CoV-2 (SC2-unique, green) or shared with other species of the Nidovirales order (CoV-common, purple) expressed as repertoire depth (a, b) and breadth (c, d). Lines represent an estimate of the central tendency of the respective values combined within each disease severity group with a 95% confidence interval shown as shadow areas when multiple data points are available at overlapping time points.

